# BNIP3L-mediated mitophagy triggered by *Brucella* in host cells is required for bacterial egress

**DOI:** 10.1101/2022.08.31.505824

**Authors:** Jérémy Verbeke, Youri Fayt, Lisa Martin, Oya Yilmaz, Jaroslaw Sedzicki, Angeline Reboul, Michel Jadot, Patricia Renard, Christoph Dehio, Henri-François Renard, Jean-Jacques Letesson, Xavier De Bolle, Thierry Arnould

## Abstract

The facultative intracellular pathogen *Brucella abortus* interacts with several organelles of the host cell to reach its replicative niche inside the endoplasmic reticulum. However, little is known about the interplay between the bacteria and the host cell mitochondria. Here, we showed that *B. abortus* triggers a strong mitochondrial network fragmentation accompanied by mitophagy and the formation of mitochondrial *Brucella*-containing vacuoles in the late steps of cellular infection. The expression of the mitophagy receptor BNIP3L induced by *B. abortus* is essential for these events and relies on the iron-dependent stabilization of the hypoxia-inducible factor 1 alpha. Functionally, BNIP3L-mediated mitophagy appears to be advantageous for bacterial exit of the host cell as BNIP3L depletion drastically reduced the number of reinfection events. Altogether, these findings highlight the intricate link between *Brucella* trafficking and the mitochondria during host cell infection.

## INTRODUCTION

Bacteria of the genus *Brucella* are Gram-negative, facultative, intracellular pathogens responsible for brucellosis, a worldwide zoonosis affecting various hosts including domestic animals (mainly cattle, sheep and goats) and humans (Atluri *et al*., 2011; González-Espinoza *et al*., 2021).*Brucella* spp. has evolved several strategies to enter, survive and proliferate inside host cells including professional phagocytes (macrophages and dendritic cells) as well as non-phagocytic cells (trophoblasts, fibroblasts and epithelial cells) (von Bargen *et al*., 2012). After its internalisation inside the host cell, *Brucella* is found within a membrane-bound vacuole, the *Brucella*-containing vacuole (BCV), which traffics along the endocytic pathway to reach an acidified compartment, the endocytic BCV (eBCV), around 8 h post-infection (pi) (Celli *et al*., 2003). This endocytic stage is necessary for the production of *Brucella* VirB Type IV secretion system (T4SS) (Boschiroli *et al*, 2002) which injects effector proteins inside the host cell (Starr *et al*., 2008). Most of the described *Brucella* effectors were shown to be required for the establishment of the replicative niche inside of the endoplasmic reticulum (ER) (Ke *et al*., 2015; Smith *et al*., 2020). Indeed, *Brucella* effectors coordinate the interaction between the eBCV and the ER exit sites (ERES) (Celli *et al*., 2005) as well as the manipulation of the Golgi vesicular trafficking (Miller *et al*., 2018; Borghesan *et al*., 2021) to generate a replicative BCV (rBCV), a compartment in continuity with the ER in which bacteria start a massive proliferation around 12 h pi (Celli *et al*., 2003; Sedzicki *et al*., 2018). In a third step, around 48 h pi, the rBCVs get engulfed inside LAMP-1-positive autophagic membranes, converting them into autophagic BCVs (aBCVs), in which *Brucella* subverts a part of the autophagy initiation complex machinery, such as ULK1, Beclin1 and ATG14L, to eventually promote bacterial egress and further reinfection of surrounding cells (Starr *et al*., 2012).

Regarding its intracellular life cycle, *Brucella* affects the functions and/or morphology of most of the host cell organelles such as lysosomes, ER and the Golgi apparatus, leading to stress and adaptive responses in the host cell (Smith *et al*., 2013; Byndloss *et al*., 2019). However, the putative interplay between *Brucella* and the mitochondrial population of infected cells is still poorly studied. The ER, lysosomes and mitochondria are able to physically and functionally interact with each other (Hamasaki *et al*., 2013; Boutry and Kim, 2021). As the central role of mitochondria in innate immunity is now well recognized (Mills *et al*., 2017; Missiroli *et al*.,2020), we hypothesised that mitochondria might be affected and play a role during *Brucella* infection. Our previous work pointed out that BCVs might physically interact with mitochondria of host cells both *in cellulo* and *in vivo* (Lobet *et al*., 2018). In addition, we showed that *B. abortus* triggers the fragmentation of the mitochondrial network in myeloid and non-myeloid cells at 48 h pi (Lobet *et al*., 2018). Mitochondrial fission plays a major role in mitochondrial quality control as it allows the segregation of damaged mitochondria upon several stresses such as calcium-mediated apoptosis (Szabadkai *et al*., 2004), oxidative stress through reactive oxygen species (ROS) (Wang *et al*., 2012) or lipopolysaccharide (LPS)-induced inflammation (Shi *et al*., 2019). Eventually, damaged mitochondrial fragments can be cleared through mitophagy, a selective autophagy process targeting mitochondria for stress-induced degradation or basal recycling through the lysosomal compartments (Onishi and Okamoto, 2021).

During mitophagy, targeted mitochondria interact with the autophagy protein LC3 inserted in autophagic phagophores, which are recruited around their cargo, leading to the formation of mitophagosomes (Kabeya *et al*., 2003; Zachari and Ktistakis, 2020). Mitophagy pathways are classified based on their dependency on ubiquitin or not. Canonical ubiquitin-dependent mitophagy mainly relies on the activity of PINK-1/Parkin. Upon loss of mitochondrial membrane potential (MMP), the PINK1 kinase is stabilised, recruits and activates the Parkin E3-ubiquitin ligase which ubiquitinates mitochondrial outer membrane proteins leading to the recruitment of receptor proteins (such as p62 or OPTN) that bind to LC3 (Agarwal and Muqit, 2022). On the other hand, ubiquitin-independent mitophagy is mediated by several mitophagy receptors such as FUNDC1, BNIP3 and BNIP3L (also called NIX), FKBP8 or Bcl2-L-13, which directly bind to LC3 (Onishi *et al*., 2021).

Here, we show, for the first time, that the mitochondrial network fragmentation induced by *B. abortus* in host cells is accompanied by an induction of mitophagy in the late steps of infection. BNIP3L, whose gene expression is regulated by HIF-1α in an iron-dependent manner during infection, is responsible for *B. abortus-induced* mitophagy. Moreover, BNIP3L-mediated mitophagy is necessary for proper *B. abortus* egress as BNIP3L depletion reduces aBCV formation and prevents reinfection. Furthermore, we also discovered a new type of BCV, that we called mitochondrial BCV (mBCV), characterised by the presence of *B. abortus* inside swollen mitochondria during the late steps of infection. Altogether, our discovery of a functional cross-talk taking place between *Brucella* and host cell mitochondria highlight what could be a crucial step for this bacterial infectious cycle.

## RESULTS

### *B. abortus* triggers mitochondrial network fragmentation

We previously showed that *B. abortus* induces the mitochondrial network fragmentation of myeloid (RAW264.7 macrophages and bone marrow-derived macrophages) and non-myeloid cells (HeLa) at 48 h pi but not at 24 h pi (Lobet *et al*., 2018). However, the kinetics of mitochondrial fragmentation, especially between 24 and 72 h pi, was not studied. To determine the precise onset of mitochondrial fragmentation induced by *B. abortus*, we analysed the morphological changes of mitochondria by assessing the aspect ratio (AR) and end-point/branched-point ratio (EBR) in infected HeLa cells displaying mitochondria immuno-stained for TOMM20 (translocase of the outer mitochondrial membrane of 20 kDa) between 8 and 72 h pi (Fig. 1A-B). While the mitochondrial network is significantly fragmented at 48 h pi, this phenotype already appears progressively from 32 h pi and is maintained until 72 h pi (Fig. 1C-D).

**Figure 1.**
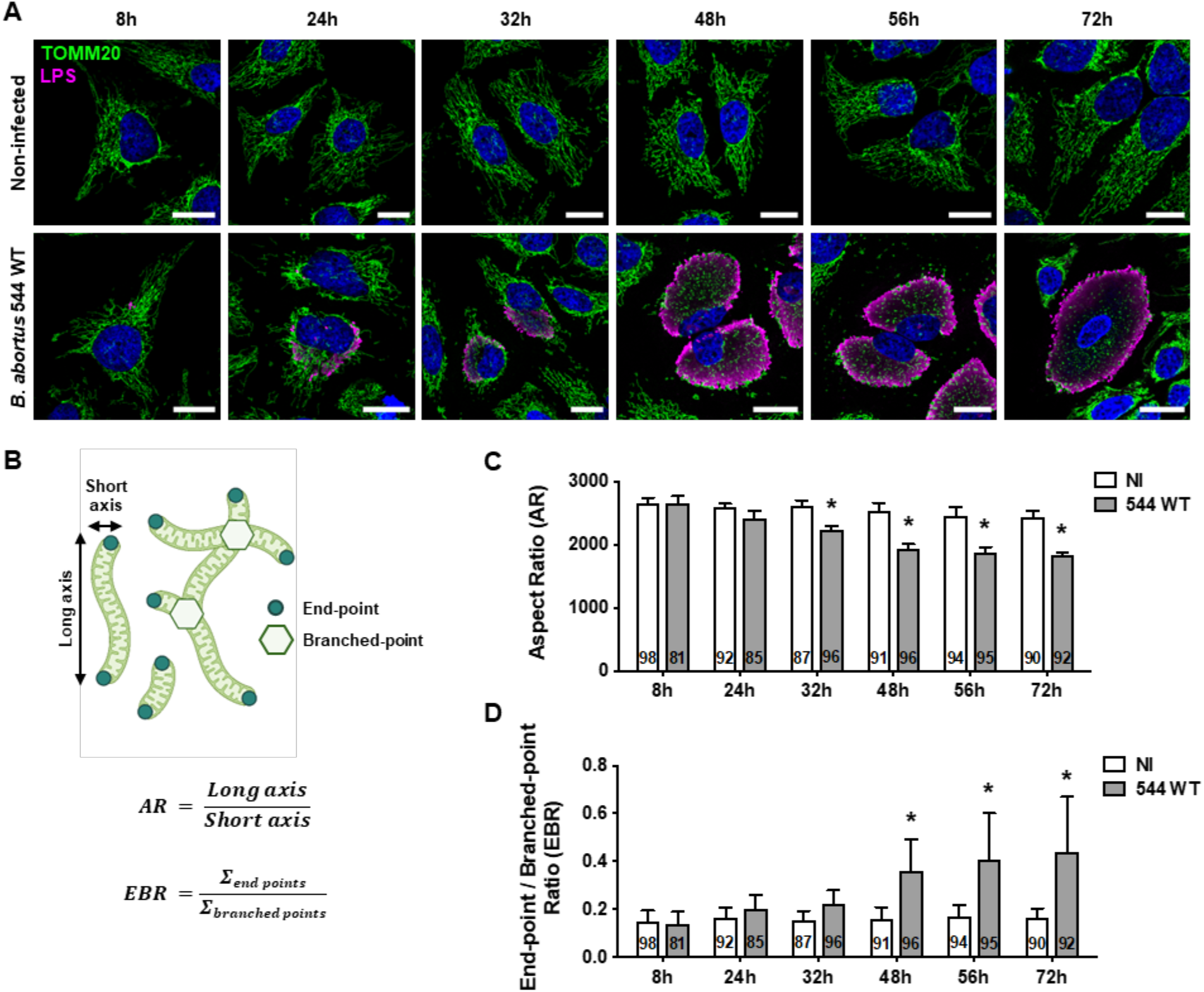
*B. abortus* triggers mitochondrial network fragmentation during the late steps of infection in HeLa cells. A. Representative confocal micrographs of HeLa cells infected or not with *B. abortus* 544 for the indicated times, then fixed and immunostained for TOMM20 (Alexa Fluor 488 – Green) and *B. abortus* LPS (Alexa Fluor 568 – Magenta). DNA was stained with Hoechst 33258 (Blue). Scale bars: 20 μm. B. Schematic summary of the calculation of the aspect ratio (AR) and the end-point/branched-point ratio (EBR) of the mitochondrial network. See the *“Quantification of mitochondrial network morphology”* paragraph in the Methods section for further information. C., D. Quantification of the mitochondrial population morphology by assessing the AR (C) and EBR (D) of the mitochondria of HeLa cells infected or not (NI) with *B. abortus* 544 for the indicated times from micrographs shown in (A). Data are presented as mean ±SD from n=5 independent experiments (the numbers indicated in the columns represent the number of cells analysed per condition). Statistical analyses were performed using a multiple Mann-Whitney test followed by a Holm-Šidàk’s multiple comparisons test; asterisks indicate significant differences compared to the control (NI); *: *p* <0.05.

### *B. abortus* is observed inside swollen mitochondria in a fraction of infected cells

Alongside the established mitochondrial network fragmentation triggered by *B. abortus* (Fig. 1), another change in mitochondrial morphology was observed between 48 and 72 h pi (Fig. 2A). Indeed, in about 10 % of infected HeLa cells, the TOMM20 signal was completely rearranged into large vesicles of a diameter of up to 5 μm. This phenotype was more frequently observed over time reaching 25 % of infected cells at 72 h pi (Fig. 2B). These TOMM20-positive structures were confirmed to be mitochondria since they colocalise with other mitochondrial markers such as the β-subunit of the ATP synthase (marker of the inner mitochondrial membrane, IMM) and the MitoTracker™ Orange (MTO) fluorescent probe which labels mitochondria in a membrane potential-dependent manner (Fig. 2C-D).

**Figure 2.**
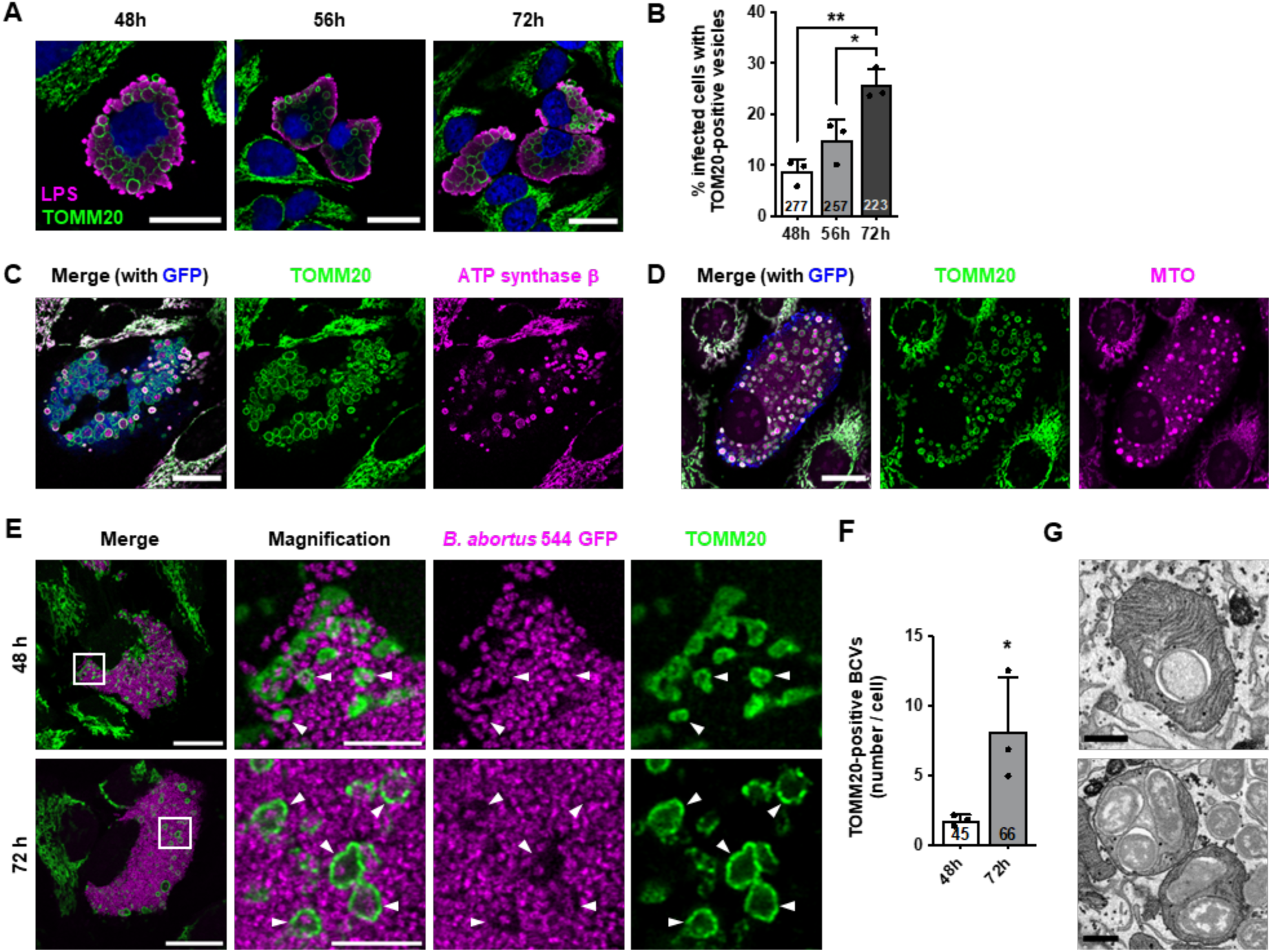
*B. abortus* is found inside swollen mitochondria of a fraction of HeLa cells during the late steps of infection. A. Representative confocal micrographs of HeLa cells infected or not with *B. abortus* 544 for the indicated times, then fixed and immunostained for TOMM20 (Alexa Fluor 488 – Green) and *B. abortus* LPS (Alexa Fluor 568 – Magenta). DNA was stained with Hoechst 33258 (Blue). Scale bars: 20 μm. B. Quantification of the percentage of infected HeLa cells displaying TOMM20-positive enlarged vesicles at the indicated times, from micrographs shown in (A). Data are presented as mean ±SD from n=3 independent experiments (the numbers indicated in the columns represent the number of cells analysed per condition); Statistical analyses were performed using a one-way ANOVA followed by a Tukey’s multiple comparisons test; *: *p* <0.05; **: *p* <0.01. C. Representative confocal micrographs of HeLa cells infected or not with *B. abortus* 544 GFP (Blue) for 48 h, then fixed and immunostained for TOMM20 (Alexa Fluor 647 – Green) and the β-subunit of the ATP synthase (Alexa Fluor 568 – Magenta). Scale bars: 20 μm. D. Representative confocal micrographs of HeLa cells infected or not with *B. abortus* 544 GFP (Blue) for 48 h, stained with 100 nM of MitoTracker™ Orange (MTO) fluorescent probe (Magenta) for 30 min before analysis, then fixed and immunostained for TOMM20 (Alexa Fluor 647 – Green). Scale bars: 20 μm. E. Representative confocal micrographs of HeLa cells infected or not with *B. abortus* 544 GFP (Magenta) for 48 and 72 h, then fixed and immunostained for TOMM20 (Alexa Fluor 647 – Green). DNA was stained with Hoechst 33258 (Blue). Arrows indicate when *B. abortus* was found inside a mitochondrion. Scale bars: 20 μm. Inset scale bars: 5 μm. F. Quantification of the number of TOMM20-positive BCVs per infected HeLa cells from micrographs shown in (E). Data are presented as mean ±SD from n=3 independent experiments (the numbers indicated in the columns represent the number of cells analysed per condition). Statistical analyses were performed using an unpaired two-tailed Student’s t-test; *: *p* = 0.05. G. FIB/SEM micrographs of HeLa cells infected with *B. abortus* 2308 RFP for 48 h. Scale bars: 600 nm.

Based on these observations, we wondered whether bacteria could be found inside of these swollen mitochondria. To address this question, we used a high resolution Airyscan microscopy and were able to detect one or few bacteria in some of these mitochondria at 48 and 72 h pi. (Fig. 2E and Movie 1). In rare cases, we even observed extreme phenotypes at 72 h pi where swollen mitochondria were full of bacteria, as depicted in stimulated emission depletion (STED) microscopy micrographs (Fig. S1). Counting of these new *Brucella*-containing compartments, that we called mitochondrial BCVs (mBCVs), showed that they are more abundant at 72 h pi (Fig. 2F). In addition, focused ion beam / scanning electron microscopy (FIB/SEM) confirmed the presence of *B. abortus* inside the mitochondrial intermembrane space (MIS) of infected cells (Fig. 2G and Movie 2).

### *B. abortus* triggers Parkin-independent mitophagy

As mitochondrial network fragmentation is a major hallmark of subsequent mitophagy, we analysed whether mitophagy was initiated in *B. abortus-infected* cells by monitoring the recruitment of phagophores at the mitochondria. To address this question, we quantified the co-localisation between LC3 and the β-subunit of the ATP synthase in infected HeLa cells (Fig. 3A and S2). A significant increase in the frequency of co-localisation events between the two proteins was observed at 48 and 72 h pi when compared to non-infected cells, while no significant difference was found at 24 h pi (Fig. 3B). In order to assess the mitophagy at a functional level, we used a FIS1-GFP-mCherry reporter construct consisting of the FIS1 (Fission 1 mitochondrial receptor) gene sequence which is tandem-tagged with the GFP and mCherry sequences. This tool allows the discrimination of acidified mitochondria (which are FIS1-mCherry-positive and GFP-negative when GFP fluorescence is quenched below pH 6 as inside of lysosomes) from neutral mitochondria (which are FIS1-GFP and -mCherry-positive) (Allen *et al*., 2013). The number of FIS1-mCherry-positive and GFP-negative structures is significantly higher in *B. abortus*-infected cells when compared to non-infected cells (Fig. 3C-D). These results confirm that *B. abortus* triggers mitophagy in infected cells.

**Figure 3.**
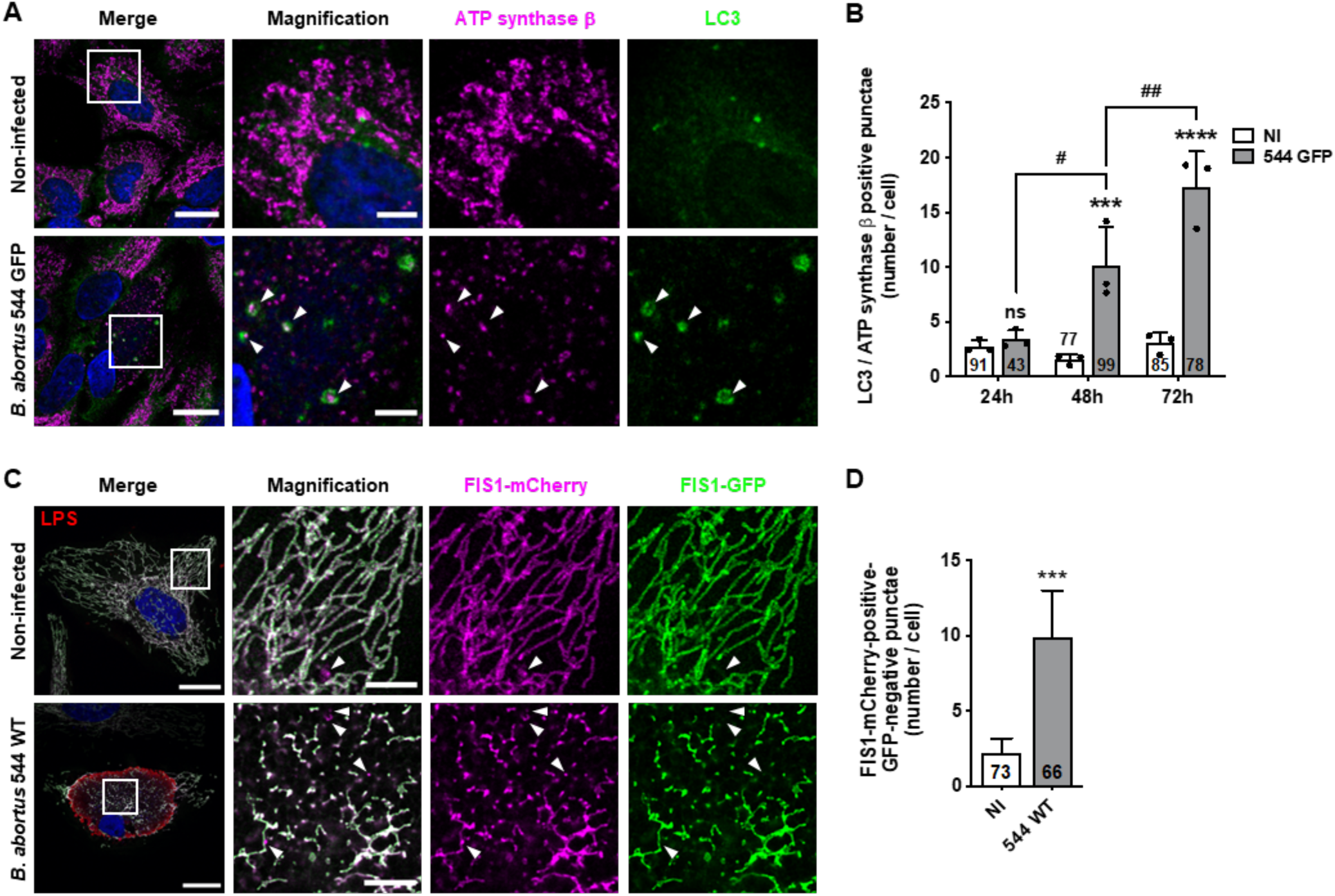
*B. abortus* triggers mitophagy in HeLa cells. A. Representative confocal micrographs of HeLa cells infected or not with *B. abortus* 544 GFP for 48 h, then fixed and immunostained for the β-subunit of the ATP synthase (Alexa Fluor 633 – Magenta) and LC3 (Alexa Fluor 568 – Green). DNA was stained with Hoechst 33258 (Blue). Arrows indicate LC3-ATP synthase β-positive punctae. Scale bars: 20 μm. Inset scale bars: 5 μm. B. Quantification of the number of LC3-β-subunit of the ATP synthase-positive punctae per HeLa cell infected or not (NI) with *B. abortus* 544 GFP for 24, 48 and 72 h from micrographs shown in (A) and (Fig EV3A-C). Data are presented as mean ±SD from n=3 independent experiments (the numbers indicated in the columns represent the number of cells analysed per condition). Statistical analyses were performed using a two-way ANOVA followed by a Šidàk’s multiple comparisons test; asterisks indicate significant differences compared to the control (NI); ns: not significant; ***: *p* <0.001; ****: *p* <0,0001; hashtags indicate significant differences between infected conditions throughout time; #: *p* <0.05; ##: *p* <0.01. C. Representative confocal micrographs of HeLa cells transfected with a FIS1-GFP(Green)-mCherry(Magenta) expression construct, infected or not with *B. abortus* 544 for 48 h, then fixed and immunostained for *B. abortus* LPS (Alexa Fluor 633 – Red). DNA was stained with Hoechst 33258 (Blue). Arrows indicate FIS1-mCherry-positive-GFP-negative punctae. Scale bars: 20 μm. Inset scale bars: 5 μm. D. Quantification of the number of FIS1-mCherry-positive-GFP-negative punctae per HeLa cell infected or not (NI) with *B. abortus* 544 for 48 h from micrographs shown in (C). Data are presented as mean ±SD from n=3 independent experiments (the numbers indicated in the columns represent the number of cells analysed per condition). Statistical analyses were performed using an unpaired two-tailed Student’s t-test; ***: *p* <0.001 (*p* = 0.0008).

Mechanisms leading to mitophagy pathways can be either ubiquitin-dependent (PINK1/Parkin pathway in response to a drop in MMP) or receptor-mediated. To identify whether the PINK1/Parkin pathway is triggered in *B. abortus*-infected cells, we analysed the MMP in infected HeLa cells using the MTO fluorescent probe, as well as the localisation of Parkin expressed in cells transfected with a Parkin-mCherry construct. *B. abortus*-infected cells did not show any loss of MMP nor Parkin co-localisation with TOMM20, as observed by confocal microscopy and flow cytometry (Fig. S3A-C). FCCP treatment was used here as a positive control since it is well-known to induce a drop in MMP, stabilise PINK1 and recruit Parkin at the surface of mitochondria which initiates mitophagy (Narendra *et al*., 2008) (Fig. S3A-C). Altogether, these observations strongly suggest that *B. abortus*-mediated mitophagy is Parkin-independent.

### *B. abortus* induces HIF-1α stabilisation and BNIP3L expression in a hypoxia-independent manner

The occurrence of some receptor-mediated mitophagy was then investigated. Since *B. abortus* is an aerobic pathogen and regarding the massive load of bacteria observed in host cells at 48 and 72 h pi, we first tested whether a hypoxic environment was generated in infected cells, by monitoring the activation of the HIF-1α/BNIP3L axis. HIF-1α is the α subunit of the HIF-1 (hypoxia inducible factor 1) transcription factor which is the master regulator of the cell response to hypoxia (Lee *et al*., 2020). BNIP3L (BCL2/adenovirus E1B 19 kDa protein-interacting protein 3-like) is a mitophagy receptor found at the OMM and the gene encoding this protein is directly regulated by HIF-1 (Daskalaki *et al*., 2018). *B. abortus-infected* cells show a nuclear localisation of HIF-1α starting at 24 h pi and reaching almost all the infected cell population at 48 and 72 h pi (Fig. 4A and B). In parallel, the abundance of BNIP3L is significantly increased in infected cells at 48 h pi when compared to non-infected cells and the protein is localised at the mitochondria as expected (Fig. 4C and D), suggesting that the HIF-1α/BNIP3L axis is activated by *B. abortus*. However, using the EF5 compound known upon hypoxic stress to form adducts which can be detected by specific antibodies (Conway *et al*., 2018), we could not detect the occurrence of hypoxic environment in infected cells, no matter the time point (Fig. S4A-C), while the EF5 compound could be abundantly detected in cells exposed to mild or severe hypoxia (Fig. S4D). These results suggest that *B. abortus* triggers HIF-1α stabilisation by a hypoxia-independent mechanism.

**Figure 4.**
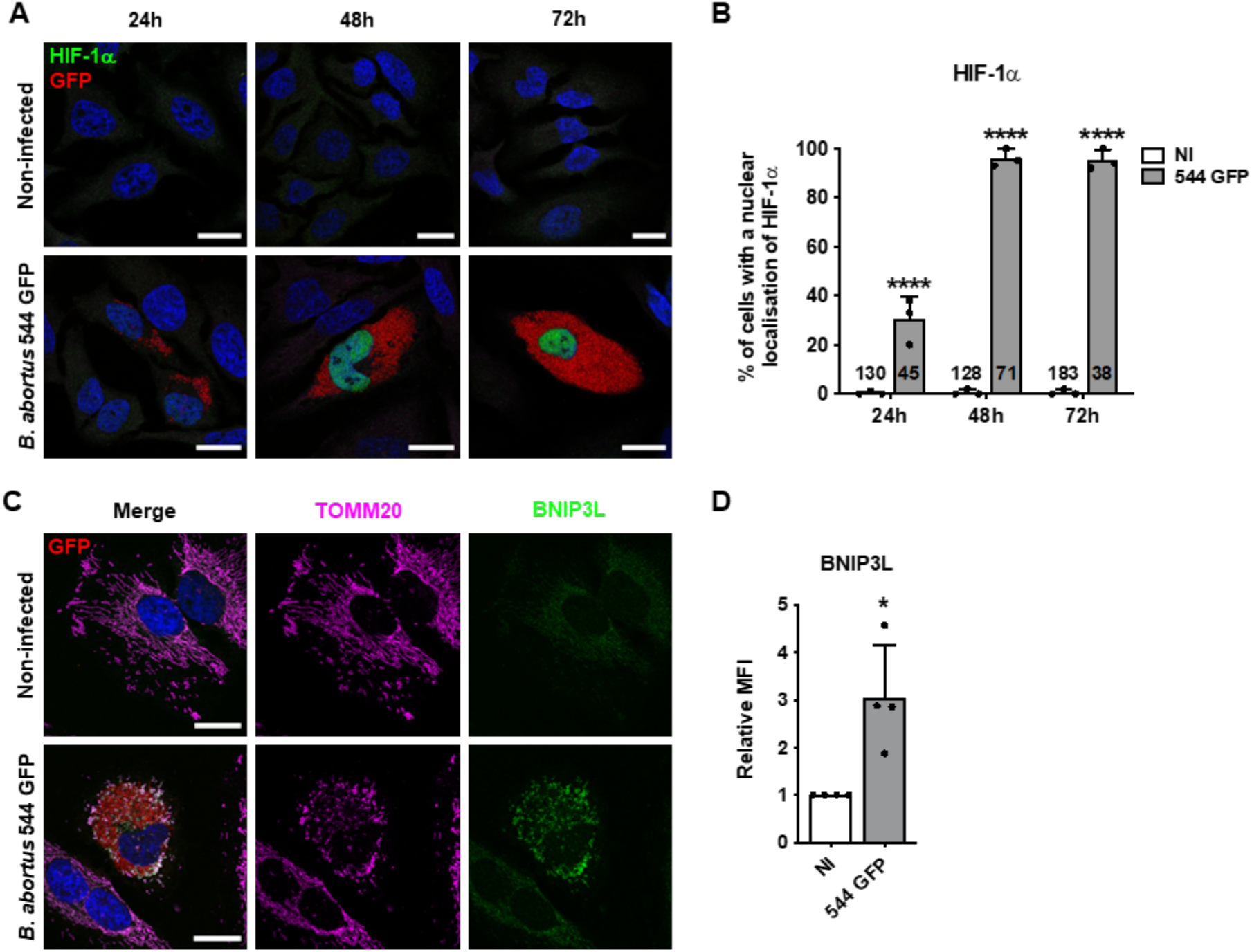
*B. abortus* induces HIF-1α stabilisation and BNIP3L expression in HeLa cells. A. Representative confocal micrographs of HeLa cells infected or not with *B. abortus* 544 GFP (Red) for 24 h, 48 h, and 72 h then fixed and immunostained for HIF-1α (Alexa 568 – Green). DNA was stained with Hoechst 33258 (Blue). Scale bars: 20 μm. B. Quantification of the percentages of cells positive for a nuclear localisation of HIF-1α from HeLa cells infected or not (NI) with *B. abortus* 544 GFP for 24, 48 and 72 h from micrographs shown in (A.B.C). Data are presented as mean ±SD from n=3 independent experiments (the numbers indicated in the columns represent the number of cells analysed per condition). Statistical analyses were performed using a two-way ANOVA followed by a Šidàk’s multiple comparisons test; asterisks indicate significant differences compared to the control (NI); ****: *p* <0.0001. C. Representative confocal micrographs of HeLa cells infected or not with *B. abortus* 544 GFP (Red) for 48 h, then fixed and immunostained for BNIP3L (Alexa 568 – Green) and TOMM20 (Alexa 633 – Magenta). DNA was stained with Hoechst 33258 (Blue). Scale bars: 20 μm. D. Relative median fluorescence intensity (MFI) of BNIP3L immunostaining from HeLa cells infected or not (NI) with *B. abortus* 544 GFP for 48 h as measured by flow cytometry. Data are presented as mean ±SD from n=4 independent experiments (8296 cells analysed in total per condition). Statistical analyses were performed using a one sample t-test; *: *p* <0.05 (*p* = 0.0351).

### Iron prevents *B. abortus-induced* HIF-1α/BNIP3L pathway activation

Regulation of HIF-1α is mainly controlled by the activity of the prolyl hydroxylase dioxygenases (PHDs) which use molecular oxygen, Fe^2+^ and alpha-ketoglutarate (αKG) to catalyse the hydroxylation of HIF-1α, leading to its proteasome-mediated degradation in normoxic conditions (Schofield and Ratcliffe, 2004). Hence, in addition to a decrease in oxygen concentration, other factors including ROS (Bell *et al*., 2007) and iron chelators such as deferoxamine (Guo *et al*., 2015) are also known to inhibit PHDs and therefore provoke HIF-1α stabilisation. As mitochondrial-derived ROS (mtROS) production is often associated with mitochondrial dysfunction, we first tested whether mtROS production is triggered upon *B. abortus* infection by using the MitoSOX™ fluorescent probe that can be oxidised by the superoxide anion radical (O_2_^-^ (Mukhopadhyay *et al*., 2007). In our experimental conditions, no increase in the MitoSOX™ fluorescence intensity was observed in *B. abortus-*infected cells when compared to non-infected cells (Fig. S5A). Moreover, neither MitoTEMPOL (a mitochondrial antioxidant) nor *N*-acetyl-L-cysteine (NAC, a cytosolic antioxidant) could prevent *B. abortus-induced* HIF-1α stabilisation and nuclear localisation (Fig. S5B-D), suggesting that HIF-1α stabilisation is not mediated by ROS.

Iron is essential for *Brucella* virulence (Roop II, 2012). A proteomic study showed that once inside the host cell, *B. abortus* remodels its iron-associated proteome through upregulation of different iron import systems, indicating that *B. abortus* faces iron starvation in the intracellular environment (Roset *et al*., 2017). We therefore tested the hypothesis that iron depletion could be responsible for *B. abortus-induced* HIF-1α stabilisation. Interestingly, iron supplementation (in the form of FeCl_2_ in the culture media) totally prevents HIF-1α nuclear localisation (Fig. 5A and B) and suppresses its stabilization (Fig. 5C), as well as the expression of BNIP3L at the mitochondria (Fig. 5D and E). These results suggest that *B. abortus* mediates HIF-1α/BNIP3L pathway activation through an iron starvation response in the host cell.

**Figure 5.**
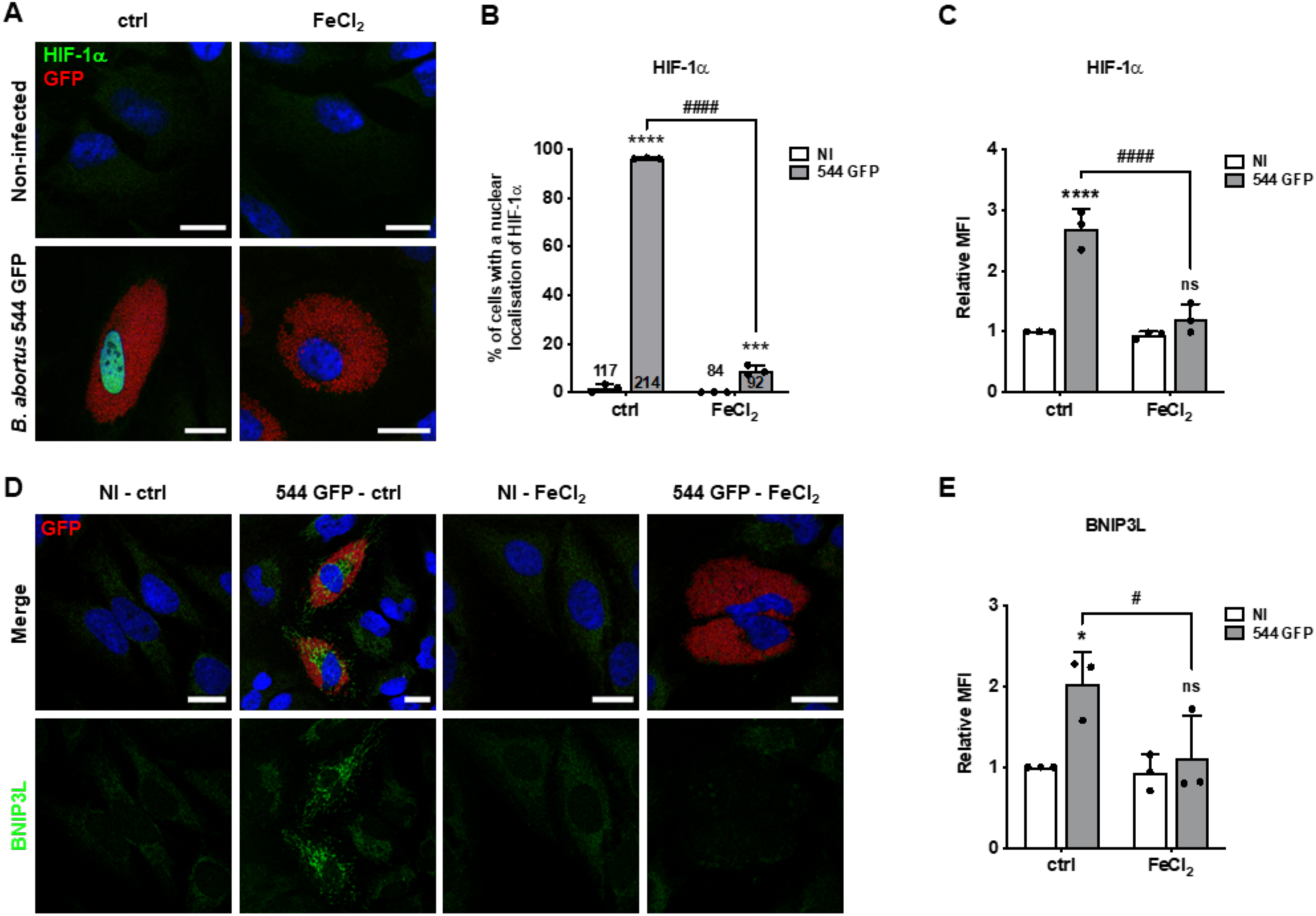
Iron supplementation prevents *B. abortus-induced* HIF-1α/BNIP3L pathway activation in HeLa cells. A. Representative confocal micrographs of HeLa cells infected or not with *B. abortus* 544 GFP (Red) treated or not (ctrl) with FeCl_2_ (500 μM) for 48 h, then fixed and immunostained for HIF-1α (Alexa 568 – Green). DNA was stained with Hoechst 33258 (Blue). Scale bars: 20 μm. B. Quantification of the percentages of cells positive for a nuclear localisation of HIF-1α from HeLa cells infected or not (NI) with *B. abortus* 544 GFP (Red) and treated or not (ctrl) with 500 μM FeCl_2_ for 48 h from micrographs shown in (A). Data are presented as mean ±SD from n=3 independent experiments (the numbers indicated in the columns represent the number of cells analysed per condition). Statistical analyses were performed using a two-way ANOVA followed by a Šidàk’s multiple comparisons test; asterisks indicate significant differences compared to the control (NI); ***: *p* <0.001; ****: *p* <0.0001; hashtags indicate significant differences compared to the infected condition without FeCl_2_; ####: *p* <0.0001. C. Relative median fluorescence intensity (MFI) of HIF-1α immunostaining from HeLa cells infected or not (NI) with *B. abortus* 544 GFP treated or not (ctrl) with 500 μM FeCl_2_ for 48 h as measured by flow cytometry. Data are presented as mean ±SD from n=3 independent experiments (10093 cells analysed in total per condition). Statistical analyses were performed using a two-way ANOVA followed by a Šidàk’s multiple comparisons test; asterisks indicate significant differences compared to the control (NI); ns: not significant; ****: *p* <0.0001; hashtags indicate significant differences compared to the infected condition without FeCl_2_; ####: *p* <0.0001. D. Representative confocal micrographs of HeLa cells infected or not with *B. abortus* 544 GFP (Red) treated or not (ctrl) with 500 μM FeCl_2_ for 48 h, then fixed and immunostained for BNIP3L (Alexa 568 – Green). DNA was stained with Hoechst 33258 (Blue). Scale bars: 20 μm. E. Relative median fluorescence intensity (MFI) of BNIP3L immunostaining from HeLa cells infected or not (NI) with *B. abortus* 544 GFP treated or not (ctrl) with 500 μM FeCl_2_ for 48 h as measured by flow cytometry. Data are presented as mean ±SD from n=3 independent experiments (10199 cells analysed in total per condition). Statistical analyses were performed using a two-way ANOVA followed by a Šidàk’s multiple comparisons test; asterisks indicate significant differences compared to the control (NI); ns: not significant; *: *p* <0,05; hashtags indicate significant differences compared to the infected condition without FeCl_2_; #: *p* <0.05.

### BNIP3L depletion prevents *B. abortus-mediated* mitophagy

To validate the putative role of BNIP3L as a mitophagy receptor induced by *B. abortus*, a gene expression silencing approach was used to knock-down the BNIP3L gene. The efficiency of BNIP3L depletion in cells transfected with a siRNA SMARTpool targeting BNIP3L mRNA on whole cell population, treated or not with CoCl_2_, known to stabilise and activate HIF-1α (Dai *et al*., 2012), was assessed by western blot analysis. The results showed that BNIP3L knock-down was very efficient at least until 72 h post-transfection when compared to cells transfected with a non-targeting control siRNA (Fig. S6). Mitochondrial network morphology was first monitored in response to BNIP3L depletion in *B. abortus-infected* cells (Fig. 6A). BNIP3L depletion significantly prevents *B. abortus-induced* mitochondrial network fragmentation. (Fig. 6B and C). Furthermore, using the FIS1-GFP-mCherry reporter construct, infected cells depleted for BNIP3L had significantly fewer FIS1-mCherry-positive and GFP-negative structures compared to infected cells transfected with a non-targeting siRNA, maintaining the number close to that of non-infected control cells (Fig. 6D and E). Altogether, these results demonstrate that BNIP3L receptor is an important actor that controls the activation of mitophagy in *B. abortus-infected* HeLa cells.

**Figure 6.**
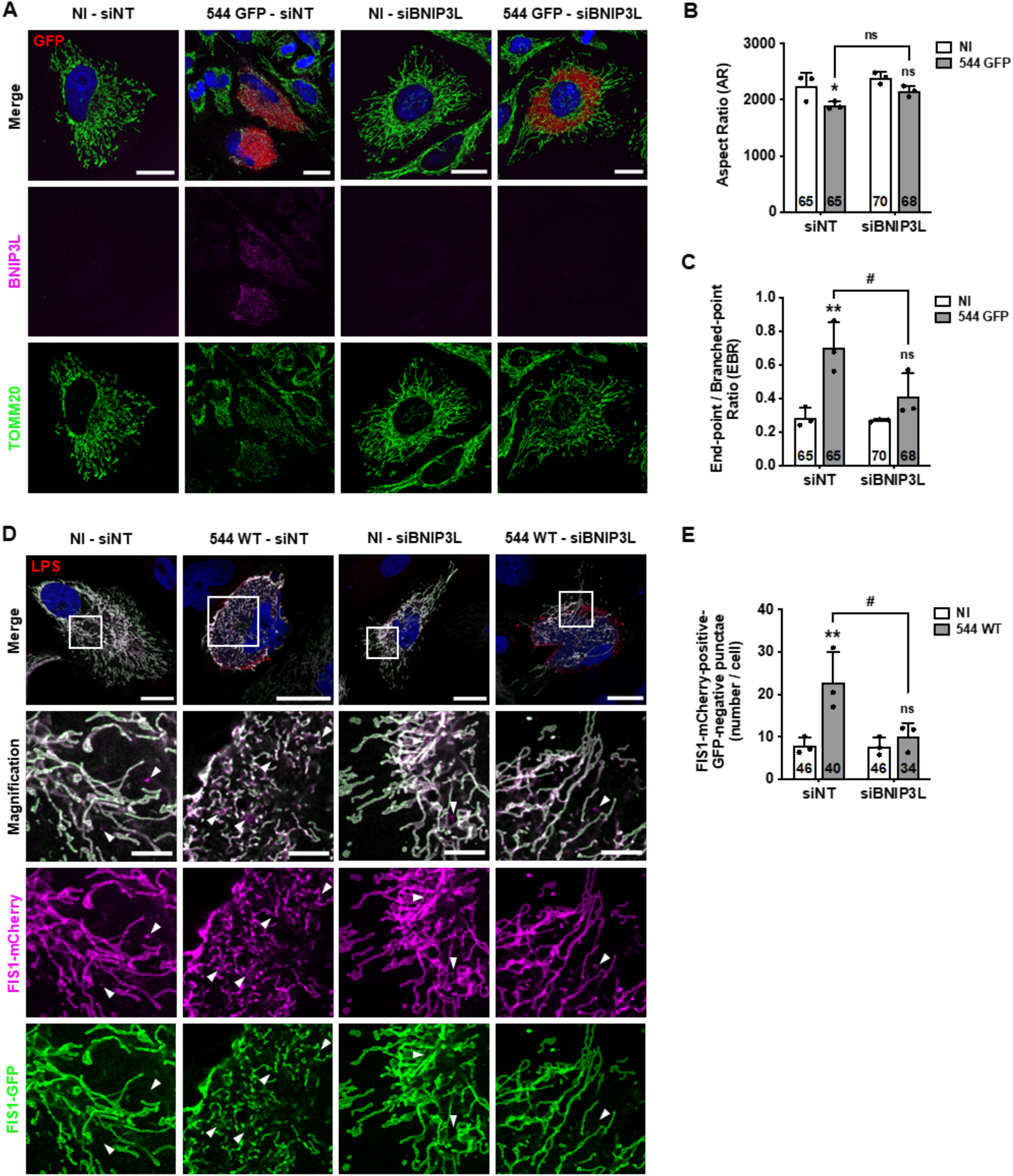
BNIP3L depletion prevents *B. abortus-induced* mitochondrial network fragmentation and mitophagy in HeLa cells. A. Representative confocal micrographs of HeLa cells transfected with a non-targeting siRNA pool (siNT – 40 nM) or a BNIP3L siRNA SMARTpool (siBNIP3L – 40 nM), infected or not (NI) with *B. abortus* 544 GFP (Red) for 48 h, then fixed and immunostained for BNIP3L (Alexa Fluor 568 – Magenta) and TOMM20 (Alexa Fluor 633 – Green). DNA was stained with Hoechst 33258 (Blue). Scale bars: 20 μm. B.,C. Quantification of the mitochondrial population morphology by assessing the AR (B) and EBR (C) of the mitochondria of HeLa cells from micrographs shown in (A). Data are presented as mean ±SD from n=3 independent experiments (the numbers indicated in the columns represent the number of cells analysed per condition). Statistical analyses were performed using a multiple Mann-Whitney test followed by a Holm-Šidàk’s multiple comparisons test; asterisks indicate significant differences compared to the control (NI); *: *p* <0.05; **: *p* <0.01; hashtags indicate significant differences compared to the infected condition transfected with a siNT; #: *p* <0.05. D. Representative confocal micrographs of HeLa cells transfected with a FIS1-GFP(Green)-mCherry(Magenta) expression construct, infected or not (NI) with *B. abortus* 544 for 48 h while being transfected with a non-targeting siRNA pool (siNT – 40 nM) or a BNIP3L siRNA SMARTpool (siBNIP3L – 40 nM), then fixed and immunostained for *B. abortus* LPS (Alexa Fluor 633 – Red). DNA was stained with Hoechst 33258 (Blue). Arrows indicate FIS1-mCherry-positive-GFP-negative punctae. Scale bars: 20 μm. Inset scale bars: 5 μm. E. Quantification of the number of FIS1-mCherry-positive-GFP-negative punctae per cell of HeLa cells from micrographs t shown in (D). Data are presented as mean ±SD from n=3 independent experiments (the numbers indicated in the columns represent the number of cells analysed per condition). Statistical analyses were performed using a two-way ANOVA followed by a Šidàk’s multiple comparisons test; asterisks indicate significant differences compared to the control (NI); ns: not significant; **: *p* <0.01; hashtags indicate significant differences compared to the infected condition transfected with a siNT; #: *p* <0.05.

### BNIP3L depletion limits aBCV formation and prevents reinfection events

Given the importance of mitochondrial function and dynamics upon bacterial infections (Spier *et al*., 2019), we next addressed the functional role of the BNIP3L-mediated mitophagy in *B. abortus* infectious cycle. Despite the fact that BNIP3L depletion does not seem to affect *B. abortus* intracellular replication (Fig. S7), an intriguing hypothesis could be that *B. abortus* egress is controlled by a BNIP3L-mediated mitophagy. Indeed, in reinfection-permissive conditions, the formation of reinfection foci is significantly decreased upon BNIP3L depletion in infected cells at 72 h pi (Fig. 7A and B). Consistent with these results, and in the same reinfection-permissive conditions, the number of bacteria found in the supernatant at 72 h pi was significantly reduced by 2-fold in BNIP3L depleted infected cells (Fig. 7C). As bacterial egress was associated with aBCV formation (Starr *et al*., 2012), we next counted the number of LAMP1-positive BCVs in infected HeLa cells depleted or not for BNIP3L. We found that the number of LAMP-1-positive BCVs is significantly reduced in cells depleted for BNIP3L to a similar extend (Fig. 7D and E). Altogether, these results indicate that BNIP3L depletion severely impairs *B. abortus* exit from the host cell by a mechanism that involves an alteration of aBCV formation.

**Figure 7.**
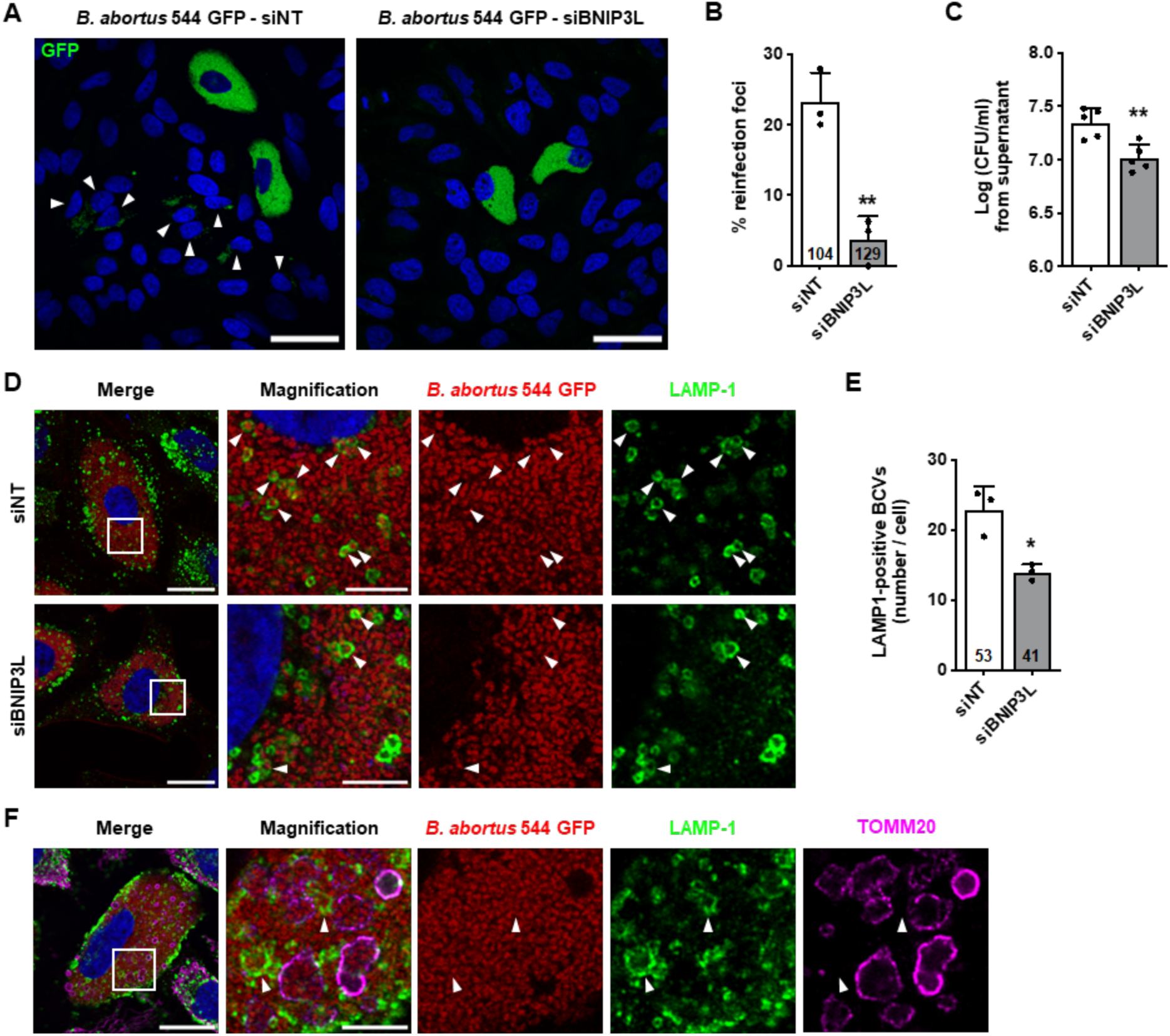
BNIP3L depletion limits aBCV formation and prevents reinfection events in HeLa cells. A. Representative confocal micrographs of HeLa cells infected with *B. abortus* 544 GFP (Green) and transfected with a nontargeting siRNA pool (siNT – 40 nM) or a BNIP3L siRNA SMARTpool (siBNIP3L – 40 nM), then incubated under reinfection-permissive conditions for 24 h before analysis at 72 h pi. Cells were fixed and DNA was stained with Hoechst 33258 (Blue). Arrows indicate reinfected cells. Scale bars: 50 μm. B. Quantification of the percentages of reinfection foci per infected cell at 72 h pi of HeLa cells from micrographs shown in (A). Data are presented as mean ±SD from n=3 independent experiments (the numbers indicated in the columns represent the number of cells analysed per condition). Statistical analyses were performed using an unpaired two-tailed Student’s t-test; **: *p* <0.01 (*p* = 0.0032). C. CFU assay expressing Log (CFU/ml) from the supernatant of HeLa cells infected with *B. abortus* 544 GFP (Green) and transfected with a non-targeting siRNA pool (siNT – 40 nM) or a BNIP3L siRNA SMARTpool (siBNIP3L – 40 nM), then incubated under reinfection-permissive conditions for 24 h before collecting the supernatant at 72 h pi for analysis. Data are presented as mean ±SD from n=5 independent experiments; Statistical analyses were performed using an unpaired twotailed Student’s t-test; (*p* = 0.0042). D. Representative confocal micrographs of HeLa cells infected with *B. abortus* 544 GFP (Red) and transfected with a nontargeting siRNA pool (siNT – 40 nM) or a BNIP3L siRNA SMARTpool (siBNIP3L – 40 nM) before analysis at 72 h pi. Cells were fixed and immunostained for LAMP-1 (Alexa Fluor 568 – Green). DNA was stained with Hoechst 33258 (Blue). Arrows indicate LAMP-1-positive BCVs. Scale bars: 20 μm. Inset scale bars: 5 μm. E. Quantification of the number of LAMP-1-positive BCVs per infected HeLa cells from micrographs shown in (E). Data are presented as mean ±SD from n=3 independent experiments (the numbers indicated in the columns represent the number of cells analysed per condition). Statistical analyses were performed using an unpaired two-tailed Student’s t-test; *: *p* <0.05 (*p* = 0.0117). F. Representative confocal micrographs of HeLa cells infected with *B. abortus* 544 GFP (Red) and transfected with a nontargeting siRNA pool (siNT – 40 nM) or a BNIP3L siRNA SMARTpool (siBNIP3L – 40 nM) before analysis at 72 h pi. Cells were fixed and immunostained for LAMP-1 (Alexa Fluor 568 – Green) and TOMM20 (Alexa Fluor 633 – Magenta). DNA was stained with Hoechst 33258 (Blue). Arrows indicate LAMP-1-positive BCVs. Scale bars: 20 μm. Inset scale bars: 5 μm.

Finally, based on the role of mitophagy in aBCV formation and the observation that the morphologies of mBCVs and aBCVs are very similar, we explored whether both structures might actually be one same type of BCV. To test this hypothesis, we performed co-immunostainings of TOMM20 and LAMP-1 in infected cells. Strikingly, our observations clearly demonstrate that TOMM20-positive BCVs are negative for LAMP-1 (Fig. 7F), which confirms that mBCVs and aBCVs are distinct organelles.

## DISCUSSION

In this study, we highlight a new mechanism demonstrating the evidence of an interplay between *Brucella* and the mitochondrial population of infected cells. We found that *B. abortus* triggers a BNIP3L-mediated mitophagy pathway which we identified to be required for the completion of its intracellular cycle up to egress and secondary infection of neighbouring cells (Fig 8). Given their function in energy production and metabolism, as well as in the regulation of inflammatory processes and programmed cell death, mitochondria are a target of choice for invading pathogens, whether they are bacteria (Lobet *et al*., 2015; Spier *et al*., 2019), viruses (Li *et al*., 2021) or protozoa (Medeiros *et al*., 2021). Even if bacteria such as *Chlamydiae spp*. inhibit mitochondrial fission to preserve the host cell ATP synthesis energy for survival and replication (Chowdhury *et al*., 2017), other pathogens such as *Listeria monocytogenes* (Carvalho *et al*., 2020), *Shigella flexneri*, (Sirianni *et al*., 2016), *Legionella pneumophila* (Escoll *et al*., 2017), and *Mycobacterium tuberculosis* (Fine-Coulson *et al*., 2015) trigger the disruption of the mitochondrial network to beneficially induce either a glycolytic shift or the host cell death. As we have shown during *B. abortus* infection, mitophagy is triggered by some of these pathogens to manipulate the host cell environment to their advantage. On the one hand, upon *L. monocytogenes* infection, a newly discovered mitophagy receptor, the NOD-like receptor NLRX1, induces mitophagy to limit mtROS production and therefore bacteria avoid to be killed by oxidant molecules (Zhang *et al*., 2019). On the other hand, in the case of *M. tuberculosis* and *M. bovis*, a transcriptomic study pointed an upregulation of BNIP3L expression in response to infection, leading to mitophagy, necessary for macrophage inflammatory function and resolution of the infection (Mahla *et al*., 2021). These two recent studies demonstrate that mitophagy can be either beneficial for the pathogen, or deleterious as part of the host cell defence.

**Figure 8.**
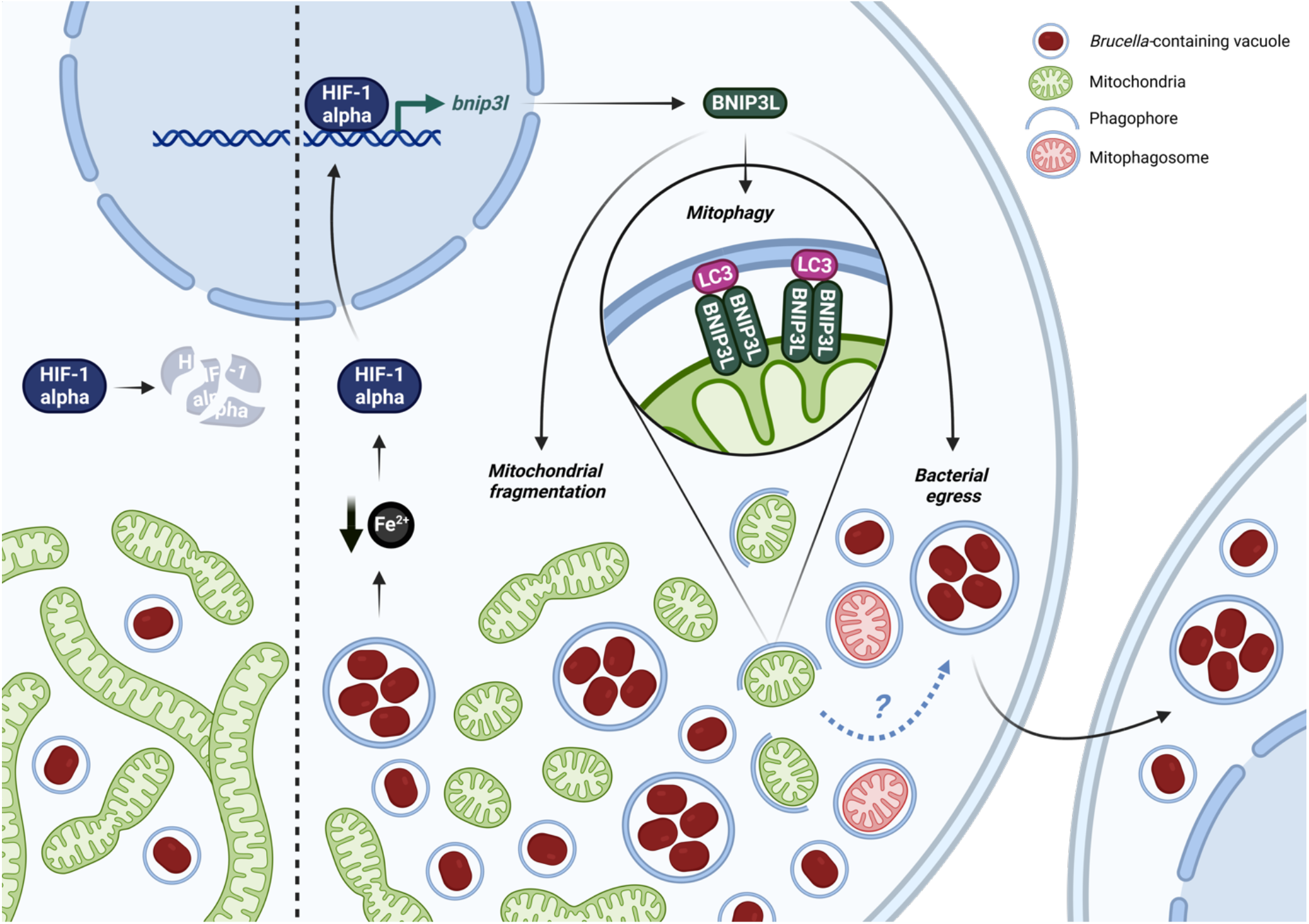
Summary of the interactions between *Brucella abortus* and the mitochondria of the host cell. In this study, we showed that until 24 h pi (A), *Brucella abortus* affects neither the mitochondrial morphology nor the endogenous degradation of HIF-1α. However, at 48 h pi (B), *B. abortus* induces an iron starvation response that stabilises and activates HIF-1α which therefore translocates to the nucleus. There, HIF-1α upregulates the expression of the mitophagy receptor BNIP3L which mediates two phenotypes during the late stages of *B. abortus* intracellular trafficking. *B.abortus*-induced BNIP3L expression mediates mitochondrial network fragmentation, one hallmark of mitophagy that is characterised by the engulfment of mitochondrial fragments inside phagophores to specifically degrade and recycle them. Eventually, BNIP3L helps to fulfil *B. abortus* intracellular cycle as it is necessary for bacterial egress and reinfection of neighbouring cells.

In the case of *B. abortus*, the induced BNIP3L-mediated mitophagy seems beneficial for the bacteria. Indeed, as BNIP3L depletion reduces the number of aBCVs in host cells, the number of bacteria in the supernatant, as well as the number of reinfected cells at 72 h pi, we demonstrated that BNIP3L is required for proper aBCV formation and bacterial egress. Our findings bring novel insights in the importance of autophagy mechanisms during the late steps of *Brucella* intracellular cycle. BNIP3L is thus a new and key actor in the list of autophagy markers such as ULK1, Beclin1 and ATG14L that are already described for aBCV formation and bacterial egress (Starr *et al*., 2012). As several organelles, including mitochondria (Hailey *et al*., 2010), have been proposed to provide membranes for autophagosome formation (Dikic and Elazar, 2018), a tempting hypothesis would be that mitochondrial turnover through mitophagy could supply membranes for aBCV formation. In addition, since mitophagy is able to protect from apoptosis through the elimination of damaged mitochondria (Szabadkai *et al*., 2004), a complementary hypothesis could be that *B. abortus* induces mitophagy to impair apoptosis of its host cell, thus keeping its replicative niche intact. This would be in agreement with the fact that *Brucella* is known to inhibit apoptosis in host cells (Ma *et al*., 2020; Ma *et al*., 2022).

Mechanistically, we showed that the induction of BNIP3L expression by *B. abortus* is dependent on the hypoxia-induced transcription factor HIF-1α (Fig. 8). The HIF-1α/BNIP3L axis has already been widely described in other mitophagy processes such as mitochondria clearance during erythrocyte maturation (Sandoval *et al*., 2008). Although mtROS-dependent HIF-1α activation has been recently reported in a macrophage model of *B. abortus* infection to be at the basis of the metabolic reprogramming induced in host cells (Gomes *et al*., 2021), no mitochondria-related HIF-1α downstream signaling was analysed. Our results strongly support that HIF-1α stabilisation and BNIP3L expression are not mediated by mtROS nor by an intracellular hypoxic environment but are dependent on iron availability. Indeed, we showed that iron supplementation was sufficient to prevent HIF-1α stabilisation and BNIP3L expression induced by *B. abortus*.These results are in line with previously reported data suggesting that a drop in iron concentrations upon *B. abortus* infection is responsible for a major rewiring of the gene expression program leading to the upregulation of different iron uptake processes, such as *Irr* (Iron-responsive regulator protein, BAB1_2175), a sensor of iron concentration variations, as well as *entC* (isochorismate synthase, BAB2_0015), one of the enzymes required for siderophore biosynthesis (Roset *et al*., 2017). Further research will be necessary to confirm the origin of iron starvation and whether it is responsible for the initiation of mitophagy. Interestingly, siderophores from the pathogenic bacteria *Pseudomonas aeruginosa* have been described to induce the stabilisation of HIF-1α, mitochondrial stress and mitophagy in *C. elegans* (Kang *et al*., 2018). Our hypothesis is that *Brucella* could consume the host cell iron through the subversion of the mitochondrial siderophore 2,5-DHBA (2,5-dihydroxybenzoic acid), a molecule with similar structure as the bacterial siderophore 2,3-DHBA (Liu *et al*., 2014), a precursor for siderophore synthesis in *Brucella* spp. (Parent *et al*., 2002), which could lead to mitochondrial stress and subsequent mitophagy.

Finally, by investigating the changes in mitochondrial morphology caused by *B. abortus* infection, we discovered a new type of BCV, the mitochondrial BCV (mBCV). Indeed, high resolution confocal and electron microscopy approaches revealed the existence of mitochondrial structures and ultra-structures – positive for several mitochondrial markers (TOMM20 and the β-subunit of the ATP synthase) – which harbour few bacteria inside the MIS between 48 and 72 h pi. The presence of bacteria inside mitochondria of host cells was also reported in the case of *Midichloria mitochondrii*, which is able to invade and colonise the MIS of oocytes in the *Ixodes ricinus* tick (Sassera *et al*., 2006; Stavru *et al*., 2020). So far, no clear underlying mechanism explaining the mitochondrial colonisation of *M. mitochondrii* has been identified yet. The authors speculate about putative fusion between *M. mitochondrii* containing vacuole and the OMM. Considering we confirmed that mBCV (LAMP-1-negative) are distinct from aBCV (LAMP-1-positive) at 72 h pi, the discovery of this new BCV highlights the diversity of potential fates of intracellular bacteria at late stages of a cellular infection.

In conclusion, our exciting findings pave the way to a better understanding of *Brucella* infection mechanisms, strengthening the intricate links existing between mitochondria of host cells and *Brucella* intracellular trafficking. Our study also emphasises that mitochondria are a central hub for invading pathogens that need to survive, proliferate and disseminate throughout the host.

## METHODS

### Reagents and antibodies

When necessary and for the indicated times, culture media were supplemented with: 20 μM carbonyl cyanide-*p*-trifluoromethoxyphenylhydrazone (FCCP; Sigma-Aldrich, C2920), 100 nM MitoTracker™ Orange CMTMRos fluorescent probe (MTO; Invitrogen, M7510), 150 μM (2-(2-nitro-1H-imidazol-1-yl)-N-(2,2,3,3,3-pentafluoropropyl) acetamide (EF5 compound; Sigma-Aldrich, EF5014), 0.5 μM MitoSOX™ fluorescent probe (Invitrogen, M36008), 10 μM MitoTEMPOL (MTEMPOL; Abcam ab144644), 5 mM *N*-acetyl-L-cysteine (NAC; Sigma-Aldrich, A9165), 500 μM iron (II) chloride tetrahydrate (FeCl_2_, Sigma-Aldrich, 44939), 100 μM cobalt chloride (CoCl_2_; Sigma-Aldrich, 15862), according to the manufacturer’s instructions.

Antibodies used for immunofluorescence and flow cytometry include: mouse anti-β subunit of the ATP synthase (1:500 dilution; Invitrogen, A21-351); rabbit anti-BNIP3L/NIX (1:100 dilution; Cell Signaling Technology, 12396S); mouse anti-*Brucella* smooth LPS (O-antigen) (A76/12G12 antibody produced as described previously in (Cloeckaert *et al*., 1990)); mouse anti-EF5-Cyanine 5 conjugate, clone ELK3-51 (1:100 dilution; Sigma-Aldrich, EF5012); rabbit anti-FAM92A (1:100 dilution; Sigma-Aldrich, HPA034760); rabbit anti-HIF-1α (1:100 dilution; Abcam, ab179483); rabbit anti-LC3B (1:100 dilution; Sigma-Aldrich, L7543); mouse anti-TOMM20 (1:100 dilution; Abcam, ab56783); rabbit anti-TOMM20 (1:200 dilution; Abcam, ab186735); Alexa Fluor 488/568/633/647 goat anti-rabbit and anti-mouse IgG conjugates (1:1000 dilution; Invitrogen); Abberior®STAR 635 goat anti-rabbit IgG conjugates (1:1000 dilution; Sigma-Aldrich). DNA was stained with Hoechst 33258 (1:5000 dilution; Invitrogen, H3569)

Antibodies used for western blot analysis include: rabbit anti-BNIP3L/NIX (1:1000 dilution; Cell Signaling Technology, 12396S); mouse anti-β actin (1:10000 dilution; Sigma-Aldrich, A5441); Ir-Dye 680/800 goat antirabbit and anti-mouse IgG conjugates (1:10000 dilution; Licor Biosciences).

### Mammalian cell culture

Human HeLa cells (HeLa CCL-2; ATCC) were cultured in Minimum Essential Medium (MEM-Glutamax; Gibco) supplemented with 1 % non-essential amino acids (100X) (MEM NEAA; Gibco), 1 mM sodium pyruvate (Gibco) and 10 % foetal bovine serum (FBS; Gibco) at 37 °C in 5 % CO_2_.

For hypoxia experiments, cells were incubated in serum-free CO_2_-independent MEM (Gibco) supplemented with 500 μM L-glutamine (Merck), and then incubated at 37 °C under either normal atmosphere (normoxia, 21 % O_2_), hypoxia (1 % O_2_), or intermediate hypoxia (between 21 and 1 % O_2_) for 3 h. To expose cells to hypoxia, a homemade pressurized incubator was used wherein gaseous N2 was injected to blow out the air contained in the incubator, thereby creating a progressive hypoxic environment. Hypoxia (1 % O_2_) and intermediate hypoxia (between 21 and 1 % O_2_) were triggered by letting air flowing out of the device for 3 min and 90 s respectively.

### Cell transfection (Plasmids and siRNA)

For Parkin localisation and mitophagy analyses respectively, a plasmid expressing Parkin-mCherry (mCherry-parkin; Addgene plasmid #23956 from R. Youle (Narendra *et al*., 2008)) and a plasmid expressing FIS1-GFP-mCherry (FIS1-GFP-mCherry; a generous gift from D. Pla-Martín (Allen *et al*., 2013)) were transfected in HeLa cells 24 h before infection using X-tremeGENE^TM^ HP DNA Transfection Reagent (Roche, 6366244001), according to the manufacturer’s instructions.

For siRNA-mediated silencing of BNIP3L, HeLa cells were transfected for 24 h before infection using DharmaFECT1 Transfection Reagent (Dharmacon, T-2001-01) with 40 nM of the ON-TARGETplus human BNIP3L siRNA SMARTpool (siBNIP3L; Dharmacon, L-011815-00-0005), according to the manufacturer’s instructions. ON-TARGETplus Non-targeting pool (siNT; Dharmacon, D-001810-10-20) was used as a control for non-specific effects. For infections with 72 h pi time points, cell transfections with siRNAs were performed 6 h after infection for 24 h in order to maintain proper BNIP3L silencing which lasts up to 72 h post-transfection (pt) (Fig. S6).

### Bacterial strains and culture

The strains used in this study were the smooth virulent *Brucella abortus* strain 544 Nal^R^ (spontaneous nalidixic acid resistant mutant strain) which is a *B. abortus* biovar 1 strain from J.-M. Verger (INRA, Tours, France) and a variant constitutively expressing the monomeric green fluorescent protein (mGFP) caused by the integration of a plasmid containing the mGFP coding sequence under the control of the *sojA* promoter (plasmid pSK *oriT kan PsojA-mgfp*, where mGFP replaces DsRed in pSK *oriT kan PsojA-dsRed* (Copin *et al*., 2012)). All *B. abortus* strains were grown in Tryptic Soy Broth-rich medium (3 % Bacto™ TSB, BD Biosciences) at 37 °C under constant agitation.

All *B. abortus* strains were handled under Biosafety level 3 (BSL-3) containment according to the Council Directive 98/81/EC of 26 October 1998, adopted by the Walloon government (July 4th, 2002).

### Cell Infection

For infections, HeLa cells were seeded either onto 13 mm glass coverslips (VWR) in 24-well plates (Corning) (for microscopy: 2×10^4^ cells/well) or in 6-well plates (Corning) (for flow cytometry: 1×10^5^ cells/well) 16 h before infection allowing cell adhesion and growth. In parallel, *B. abortus* cultures were incubated for 16 h in TSB medium from isolated colonies. The next day, bacterial cultures were washed twice in phosphate buffer saline (PBS, Lonza) and bacterial exponential growth was controlled by measuring culture optical density at 600 nm. *B. abortus* infection doses were prepared in complete HeLa culture medium at a multiplicity of infection (MOI) of 2000 and incubated with the HeLa cells. Infected cells were centrifuged at 400 g for 10 min to favour bacteria-to-host cell contact and incubated for 1 h at 37 °C in 5 % CO_2_. Cells were then incubated with 50 μg/ml of gentamycin (Gibco) for 1 h to kill the remaining extracellular bacteria and thereafter washed and incubated with 10 μg/ml of gentamycin for the desired time points.

For reinfection experiments, infected cells were incubated in so-called reinfection-permissive conditions at 48 h pi in culture media without gentamycin to allow the survival of released bacteria and reinfection of new host cells. Cells were washed once in PBS and incubated with gentamycin-free HeLa culture media until 72 h pi for further analysis.

### CFU assay

To assess *B. abortus* intracellular replication at several time points (2 h, 6 h, 24 h and 48 h pi), infected cells were washed once in PBS then lysed with 0.1 % Triton-X-100 (Sigma-Aldrich) in PBS for 10 min at room temperature. Serial supernatant dilutions were then plated onto TSB agar plates, incubated at 37 °C for 5 days and colony-forming units (CFUs) were counted for each condition from three technical replicates (Roba *et al*.,2022).

To assess *B. abortus* egress at 72 h pi, serial dilutions of supernatant from infected cells previously incubated in reinfection-permissive conditions were plated onto TSB agar plates, incubated at 37°C for 5 days and colony-forming units (CFU) were counted for each condition from three technical replicates.

### Immunofluorescence

At the different time points indicated in figure legends, cells seeded on coverslips were washed once with PBS, fixed with 4 % paraformaldehyde (PFA; VWR) for 30 min at 37 °C, then washed three times with PBS and permeabilised with 1% Triton-X-100 (Carl Roth) in PBS for 10 min. For LC3 immunostainings, fixation and permeabilisation was performed using pure ice-cold methanol (−20 °C) for 15 min at 4 °C. Cells were then incubated with a blocking solution (2 % bovine serum albumin (BSA; VWR) diluted in PBS) for 30 min at room temperature, and then incubated with the primary antibodies in blocking solution for 2 h at room temperature. Afterwards, cells were washed three times with the blocking solution and incubated with the secondary antibodies in blocking solution for 1 h a room temperature. For LAMP-1 immunostainings, permeabilisation was performed using 0.2 % saponin (Sigma-Aldrich) diluted in PBS for 30 min, and the blocking solution was composed of 0.2 % saponin, 2 % BSA, 10 % horse serum (HS, Gibco) diluted in PBS. DNA staining using Hoechst 33258 was performed during the incubation with the secondary antibody. Finally, cells were washed twice with blocking buffer and twice with distilled water, then mounted on glass slides in Mowiol (Sigma-Aldrich) or Fluoromount-G (Invitrogen™) as appropriate.

### Light microscopy

Fixed and immunostained cells were visualised with a Leica TCS SP5 II confocal laser-scanning microscope equipped with a HCX Plan Apo CS 40x numerical aperture (NA) 1.3 oil immersion objective. For observations requiring a super resolution, a Zeiss LSM 900 confocal laser-scanning microscope equipped with an Airyscan 2 multiplex system and a Plan Apo 63x NA 1.4 oil immersion objective was used, in particular for the analysis of *B. abortus* localisation inside of mitochondria and mBCVs (Fig. 2E and S1), mitophagy (Fig. 3C and 6D), reinfection foci (Fig. 7A) and aBCVs (Fig. 7D and 7F). For experiments assessing reinfection foci, the same microscope with a Plan Apo 40x NA 1.3 oil immersion objective was used instead). In addition, stimulated emission depletion (STED) microscopy analyses using an Abberior STEDycon attached to a Zeiss Axio Imager Z2 microscope were performed to better observe *B. abortus* localisation inside of mitochondria (Fig. S1)

For live cell imaging, observations were made at 37 °C with a Nikon Eclipse Ti2 inverted epifluorescence microscope equipped with a Plan Apo *λ* DM 100x K 1.45/0.13 PH3 phase-contrast objective to visualise the MitoSOX™ probe staining. Look-up tables (LUT) were adjusted to the best signal–background ratio.

### Quantitative analyses of fluorescence micrographs

All fluorescence images were analysed using FIJI v.2.1.0, a distribution of ImageJ.

#### Quantification of mitochondrial network morphology

The length and branching status of mitochondrial network were determined by calculating the aspect ratio (AR) and the end-point/branched-point ratio (EBR) of mitochondrial particles in entire cell sections as previously described (De Vos and Sheetz, 2007; Lobet *et al*., 2018). The AR represents the mean of the ratio between the long axis and the short axis of each mitochondrial fragments of one cell. The AR is therefore proportional to the length of mitochondrial fragments. The EBR represents the mean of the ratio between the end-points and branched-points of each mitochondrial fragments of one cell. The EBR is therefore proportional to the disconnected status of the mitochondrial network. (Fig. 1B). The analyses were performed on samples from 5 (n=5: Fig. 1) and 3 (n=3; Fig. 6) independent biological experiments.

#### Quantification of the percentages of TOMM20-vesicles-positive infected cells

The percentage of TOMM20-vesicles-positive infected cells was determined by calculating the ratio between the number of infected cells displaying large TOMM20-positive vesicles and the total number of infected cells from one experiment. The analyses were performed on samples from 3 independent biological experiments (n=3).

#### Quantification of LC3 recruitment at the mitochondria proximity

The quantification of LC3 recruitment at the proximity of mitochondria was determined by analysing the colocalisation between Alexa Fluor 568 (LC3) and Alexa Fluor 633 (β-subunit of ATP synthase) related fluorescence signals. A signal threshold was thus applied on Alexa Fluor 568 micrographs to dissociate LC3-type II punctae structures from LC3-type I diffuse signal, as well as on Alexa Fluor 633 micrographs in order to delineate the mitochondrial network. The number of co-localisation events between LC3 punctae and mitochondria was performed by generating on overlay of both processed images and counting the number of LC3-β-subunit of ATP synthase-positive co-localised events in entire cell sections. The analyses were performed on samples from 3 independent biological experiments (n=3).

#### Quantification of the number of mitochondrial fragments in acidified compartments

The number of mitochondrial fragments in acidified structures was determined by counting the number of FIS1-mCherry(magenta)-positive but GFP(green)-negative structures in entire cell sections as previously described (Allen *et al*., 2013). The analyses were performed on samples from 5 (n=5: Fig. 3) and 3 (n=3; Fig. 6) independent biological experiments.

#### Quantification of the percentages of cells with a HIF-1α nuclear pattern

The percentages of cells with a HIF-1α nuclear pattern were determined by calculating the ratio between the number of cells that display a nuclear localisation of HIF-1α and the total number of cells observed from one experiment. The analyses were performed on samples from 3 independent biological experiments (n=3: Fig. 4 and Fig. 5).

#### Quantification of the percentages of reinfection events

The percentages of reinfection events were determined by calculating the percentages of reinfection foci as described previously (Starr *et al*., 2012). Reinfection foci were defined as a primary infected cell (containing a massive number of bacteria – 72 h pi) surrounded by at least 4 adjacent cells containing a small number of bacteria (secondary infection – 24 h pi). The analyses were performed on samples from 3 independent biological experiments (n=3).

#### Quantification of the number of aBCVs and mBCVs

The number of aBCVs was determined by counting the number of LAMP-1-positive vesicles harbouring 1 or more GFP-positive *B. abortus* 544 bacteria in entire cell sections. The same procedure was applied to mBCVs by counting TOMM20-positive BCVs. The analyses were performed on samples from 3 independent biological experiments (n=3).

### Electron microscopy sample preparation

HeLa cells were seeded onto 32 mm gridded glass coverslips (Ibidi, Martinsried, Germany) in a 6-well plate at 150,000 cells per well and incubated for 16 h before infection. In parallel, *B. abortus* 2308 were grown overnight in TSB medium at 37 °C to an OD of 0.8-1.0. Bacteria were then diluted in DMEM / 10 % FCS and added to HeLa cells at a final MOI of 2000. Plates were centrifuged at 400 x *g* for 20 min at 4 °C to synchronize bacterial entry. After 2 h of incubation at 37 °C and 5 % CO_2_, extracellular bacteria were killed by exchanging the infection medium by DMEM / 10% FCS supplemented with 100 μg/ml gentamicin. After the total infection time, cells were fixed using PHEM fixation buffer (4 % formaldehyde, 0.2 % glutaraldehyde, 60 mM PIPES, 25 mM HEPES, 10 mM EGTA, 4 mM MgCl_2_) for 90 min. at room temperature. Following fixation, the coverslips were washed in PHEM buffer (60 mM PIPES, 25 mM HEPES, 10 mM EGTA, 4 mM MgCl_2_) and fixed in cacodylate fixation buffer (2.5 % glutaraldehyde, 150 mM sodium cacodylate, 2 mM MgCl_2_) at 4 °C overnight. Following overnight fixation, samples were washed 3 times with cacodylate buffer (150 mM sodium cacodylate, 2 mM MgCl_2_) at 4 °C; and then immersed in freshly prepared reduced osmium buffer (2 % osmium tetroxide, 150 mM sodium cacodylate, 2 mM MgCl_2_, 40 mM potassium ferrocyanide) for 1 h at 4 °C. After this initial staining/fixation step, the samples were washed with deionized water at room temperature and immersed in 100 mM thiocarbohydrazide solution for 20 min at room temperature, and then washed with deionized water and incubated in 2 % osmium tetroxide for 30 min at room temperature. This was followed by overnight incubation in 1 % uranyl acetate at 4 °C. The following morning, the samples were washed in deionized water and incubated in freshly prepared 20mM lead aspartate solution for 30 min at 60°C. Samples were then dehydrated with ethanol and immersed in 50 % solution of durcupan in ethanol for 1 h. Afterwards, the samples were incubated 2 times in fresh durcupan and placed at 60 °C for 48 h for polymerization.

### Focused ion beam / scanning electron microscopy (FIB/SEM)

The cells of interest were located in the polymerized resin block, trimmed and attached to pre-tilt 45° SEM stubs (Agar Scientific, Stansted, UK) using colloidal silver paint (Ted Pella, Redding, CA), sputter-coated with platinum and subjected to FIB/SEM tomography. The images were acquired with a Helios NanoLab 650 Dual Beam FIB/SEM using the Slice and View software (FEI, Hillsboro, OR). They had 3072 x 2048 or 2048 x 1780 pixel and were collected using an Elstar in-lens BSE detector at 1.5 kV with a horizontal field width of 15 μm at a working distance of 4.01 mm. The milling was performed with a FIB operating at 30 kV and 0.78 nA beam current. The thickness of the slices was between 10 and 20 nm. Image stacks were aligned using the TrackEM2 plugin for ImageJ (Cardona et al., 2010).

### Flow cytometry

At the different time points indicated in figure legends, cells were washed once with PBS, detached with Trypsin 0.05 % - EDTA (Gibco) for 3 min at 37 °C and centrifuged at 400 g for 5 min at 4 °C. Collected cells were washed twice with an ice-cold Flow Cytometry buffer (0.5 % BSA, 2 mM EDTA diluted in PBS) and then fixed with the IC Intracellular Fixation Buffer (eBioscience™ Invitrogen, 00-8222-49) for 30 min at room temperature. Fixed cells were permeabilised with the Permeabilisation Buffer 1X (eBioscience™ Invitrogen, 00-8333-56) for 5 min at room temperature, then incubated with the primary antibodies in Permeabilisation Buffer for 1 h at room temperature. Cells were next washed once with Permeabilisation Buffer and incubated with the secondary antibodies in Permeabilisation Buffer for 1 h a room temperature. Finally, cells were washed once with Permeabilisation Buffer and transferred into glass tubes with Flow Cytometry buffer before analysis with the BD Biosciences FACSVerse™. Data analyses were performed using the FlowJo™ software (BD Biosciences).

### Western blot analysis

At the different time points indicated in figure legend, cells were washed once with ice-cold PBS and lysed in radioimmunoprecipitated assay (RIPA) buffer (150 mM NaCl, 1 % NP40, 0.1 % SDS, 1 % sodium deoxycholate, 25 mM Tris-HCL; pH 7.6) supplemented with complete Protease Inhibitor Cocktail (Roche, 11697498001) and 4 % Phosphatase Inhibitor Buffer (25 mM Na3VO4, 250 mM 4-nitrophenylphosphate, 250 mM β-glycerophosphate, 125 mM NaF) for 10 min on ice. Cell lysates were centrifuged at 13000 g for 10 min at 4 °C. Clear cell lysates were then incubated for 1 h at 80 °C to kill remaining bacteria. Protein concentration was assessed with the Pierce 660 nm Protein Assay Reagent (Thermo Scientific, 22660). Samples of 10 μg of cell lysate proteins were resolved by gel electrophoresis using a 10 % acrylamide gel. Proteins were then electrotransferred onto a polyvinylidene fluoride (PVDF) membrane (0.45 μm) (Merck by ice-cold liquid transfer for 2 h at 70 V. Membranes were then incubated with Licor Intercept (PBS) Blocking buffer (LICOR Biosciences, 927-70003) diluted twice in PBS for 1 h at room temperature, and then incubated with the primary antibody diluted in Intercept Blocking buffer supplemented with 0.1 % Tween20 (Carl Roth, 9127.1) and 0.2 % sodium dodecyl sulfate (SDS; VWR, A3942) for 16 h at 4° C. Membranes were then washed three times with 0.1 % Tween20 diluted in PBS, and then incubated with the secondary antibody diluted in Intercept Blocking buffer supplemented with 0.1 % Tween20 and 0.2 % SDS for 1 h at room temperature. Membranes were then washed twice with 0.1 % Tween20 in PBS, once in PBS, and the fluorescence intensity of the bands corresponding to the proteins of interest were detected using the Odyssey ODY-1869 scanner (LICOR). The immunodetection of the β-actin was used as a loading control.

### Statistical analyses

Statistical analyses were performed using GraphPad Prism 9 software. All data are presented as the means ±standard deviation (SD) of results from at least three biological independent experiments (n=3). Normality of the distribution was assessed using Shapiro-Wilk tests. Comparisons between two conditions were assessed using an unpaired two-tailed Student’s t-test, or a one sample t-test when normality failed. Comparisons between more than two groups involving one single factor were assessed using a one-way ANOVA followed by a Tukey’s multiple comparisons test. Comparisons between more than two groups involving two simultaneous factors were assessed using a two-way ANOVA followed by a Šidàk’s multiple comparisons test, or a multiple Mann-Whitney test followed by a Holm-Šidàk’s multiple comparisons test when normality failed. A p-value < 0.05 was considered as statistically significant.

## ACKNOWLEDGEMENTS

The authors would like to thank Dr. Suzana Salcedo (University of Lyon, France) for her helpful discussions. The authors also thank the “Morphology and Imaging” (Morph-Im) technological platform (University of Namur) and especially Catherine Demazy and Noëlle Ninane for the advice and trainings for the use of the Leica TCS SP5 II confocal microscope. We also thank Dr. David Pla-Martín for generously providing the FIS1-GFP-mCherry reporter construct. Jérémy Verbeke is a Research Fellow (2018-2022) of the F.R.S-FNRS (Fonds de la Recherche Scientifique, Belgium). This work was also supported by two “Crédit de Recherche” grants (CDR 2019-2021: “MITOCHOBRU” grant J.0003.20-AID 35252856 and CDR 2022-2023: “Brucella and BNIP3L-mediated mitophagy” grant J.0003.22 AID 40007965) obtained from the F.R.S-FNRS, as well as by a Start-Up grant obtained from the Fondation Francqui (to H.-F.R.), and the grant 310030B_201273 obtained from the Swiss National Science Foundation (to C.D.).

## AUTHOR CONTRIBUTIONS

Conceptualization: J.V., A.R., M.J., P.R., H.-F.R., J.-J.L., X.D.B, T.A.

Investigation: J.V., Y.F., L.M., O.Y., J.S.

Formal Analysis: J.V., Y.F., L.M., O.Y., J.S.

Resources: J.S., C.D.

Writing – Original Draft: J.V., X.D.B., T.A.

Writing – Review & Editing: J.V., A.R., J.S., M.J., P.R., C.D., H.-F.R., J.-J.L., X.D.B., T.A.

Funding acquisition: J.V., C.D., H.-F.R., X.D.B., T.A.

Supervision: X.D.B., T.A.

## MOVIE LEGENDS

**Movie 1. *B. abortus* is found inside of some mitochondria of HeLa cells at 48 h pi.**

Airyscan microscopy Z-stack from HeLa cells infected with *B. abortus* 544 GFP (Magenta) for 48 h, then fixed and immunostained for TOMM20 (Alexa Fluor 647 – Green).

**Movie 2. *B. abortus* is found in the MIS of several mitochondria of HeLa cells at 48 h pi.**

FIB/SEM stack from HeLa cells infected with *B. abortus* 2308 RFP for 48 h.

## SUPPLEMENTARY FIGURES

**Figure S1.**
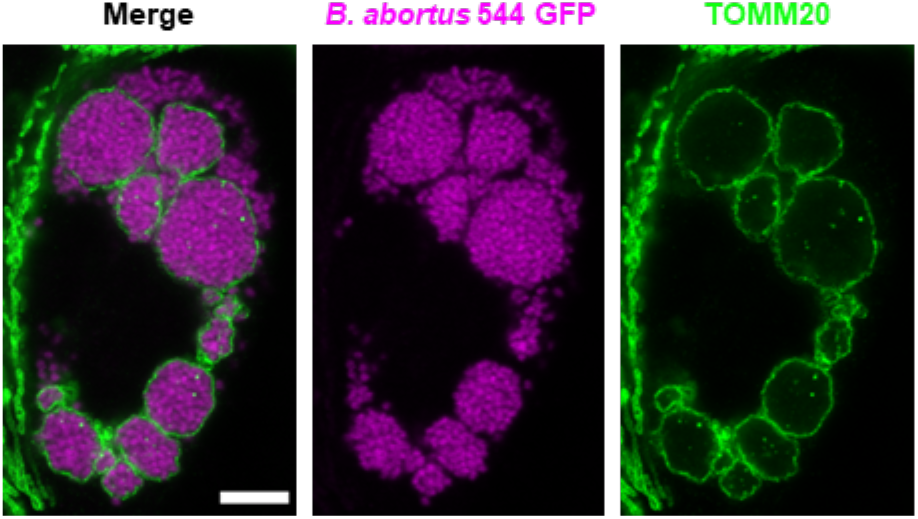
A massive load of *B. abortus* is found inside swollen mitochondria in a fraction of HeLa cells at 72 h pi. STED micrographs of a HeLa cell infected with *B. abortus* 544 GFP (Magenta) for 72 h, then fixed and immunostained for TOMM20 (Abberior^®^ STAR 635 – Green). Scale bars: 5 μm.

**Figure S2.**
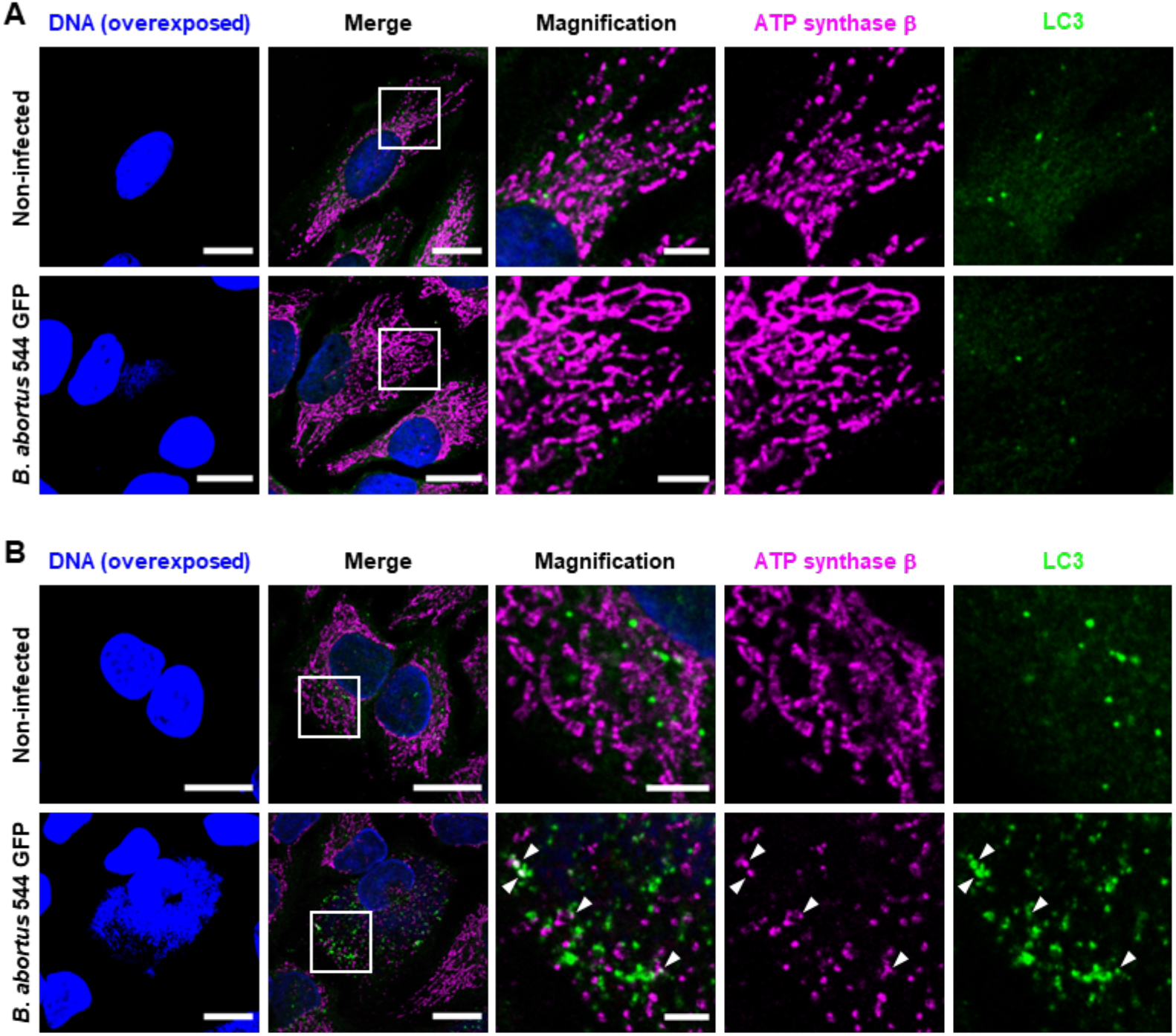
Kinetics of mitophagy triggered by *B. abortus* in HeLa cells at 24 and 72 h pi. A., B. Representative confocal micrographs of HeLa cells infected or not with *B. abortus* 544 GFP for 24 h (A.) and 72 h (B.), then fixed and immunostained for the β-subunit of the ATP synthase (Alexa Fluor 633 – Magenta) and LC3 (Alexa Fluor 568 – Green). DNA was stained with Hoechst 33258 (Blue). The Hoechst intensity was intentionally shown with overexposed signals to visualise bacterial DNA and confirm the infected cells status. Scale bars: 20 μm. Inset scale bars: 5 μm.

**Figure S3.**
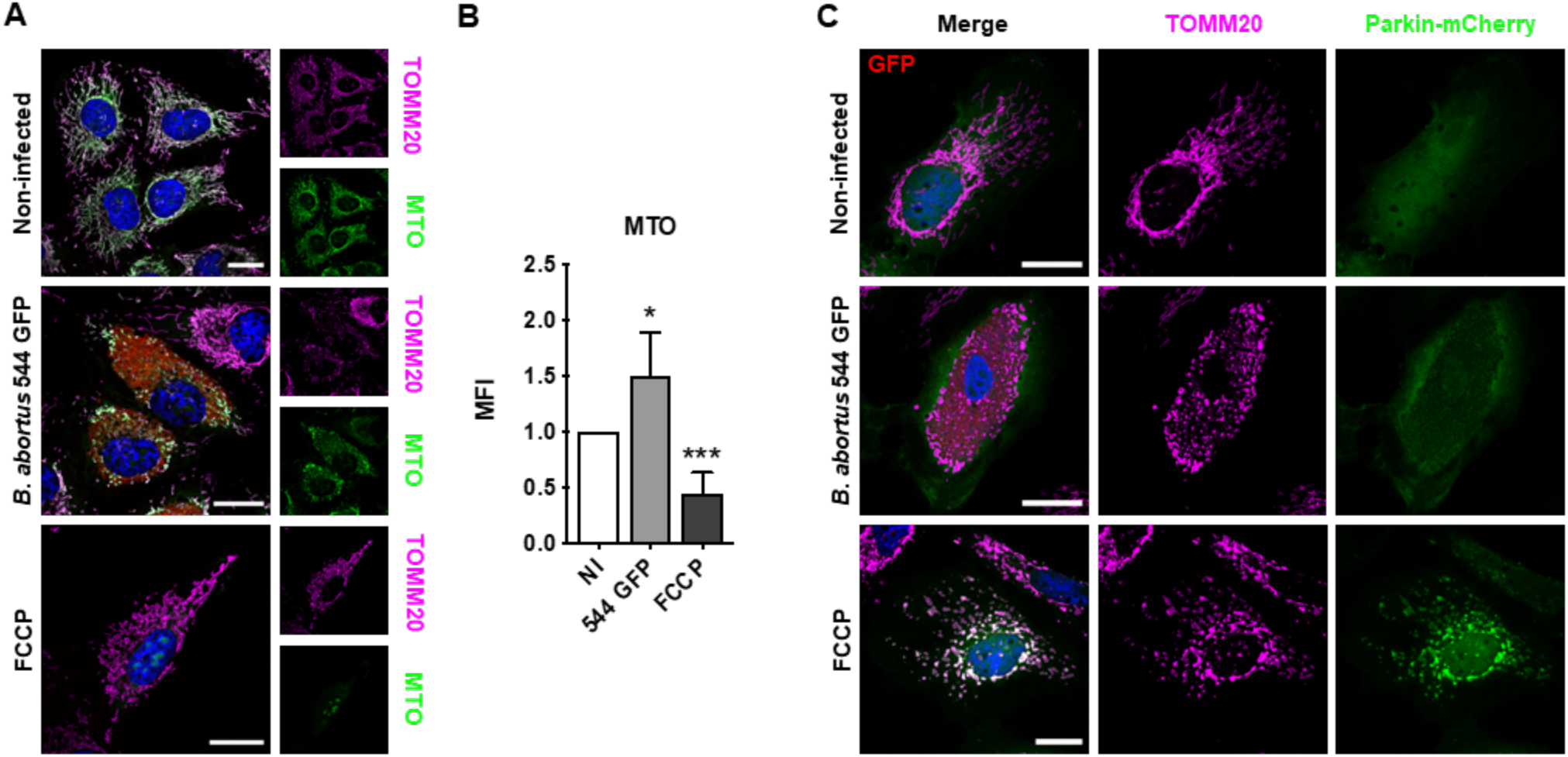
*B. abortus* triggers Parkin-independent mitophagy. A. Representative confocal micrographs of HeLa cells infected or not with *B. abortus* 544 GFP (Red) for 48 h, stained with 100 nM of MTO fluorescent probe (Green) for 30 min before analysis, then fixed and immunostained for TOMM20 (Alexa Fluor 633 – Magenta). DNA was stained with Hoechst 33258 (Blue). HeLa cells treated with 20 μM FCCP for 30 min were used as a positive control. Scale bars: 20 μm. B. Relative median fluorescence intensity (MFI) of the MTO fluorescent probe of HeLa cells infected or not (NI) with *B. abortus* 544 GFP for 48 h as measured by flow cytometry. HeLa cells treated with 20 μM FCCP for 30 min were used as a positive control. Data are presented as mean ±SD from n=5 independent experiments (9718 cells analysed in total per condition). Statistical analyses were performed using a one-way ANOVA followed by a Tukey’s multiple comparisons test; asterisks indicate significant differences compared to the control (NI); *: *p* <0.05; ***: *p* <0.001. C. Representative confocal micrographs of HeLa cells transfected with a Parkin-mCherry (Green) expression construct, infected or not with *B. abortus* 544 GFP (Red) for 48 h, then fixed and immunostained for TOMM20 (Alexa Fluor 647 – Magenta). DNA was stained with Hoechst 33258 (Blue). HeLa cells treated with FCCP (20 μM for 30 min) were used as a positive control. Scale bars: 20 μm.

**Figure S4.**
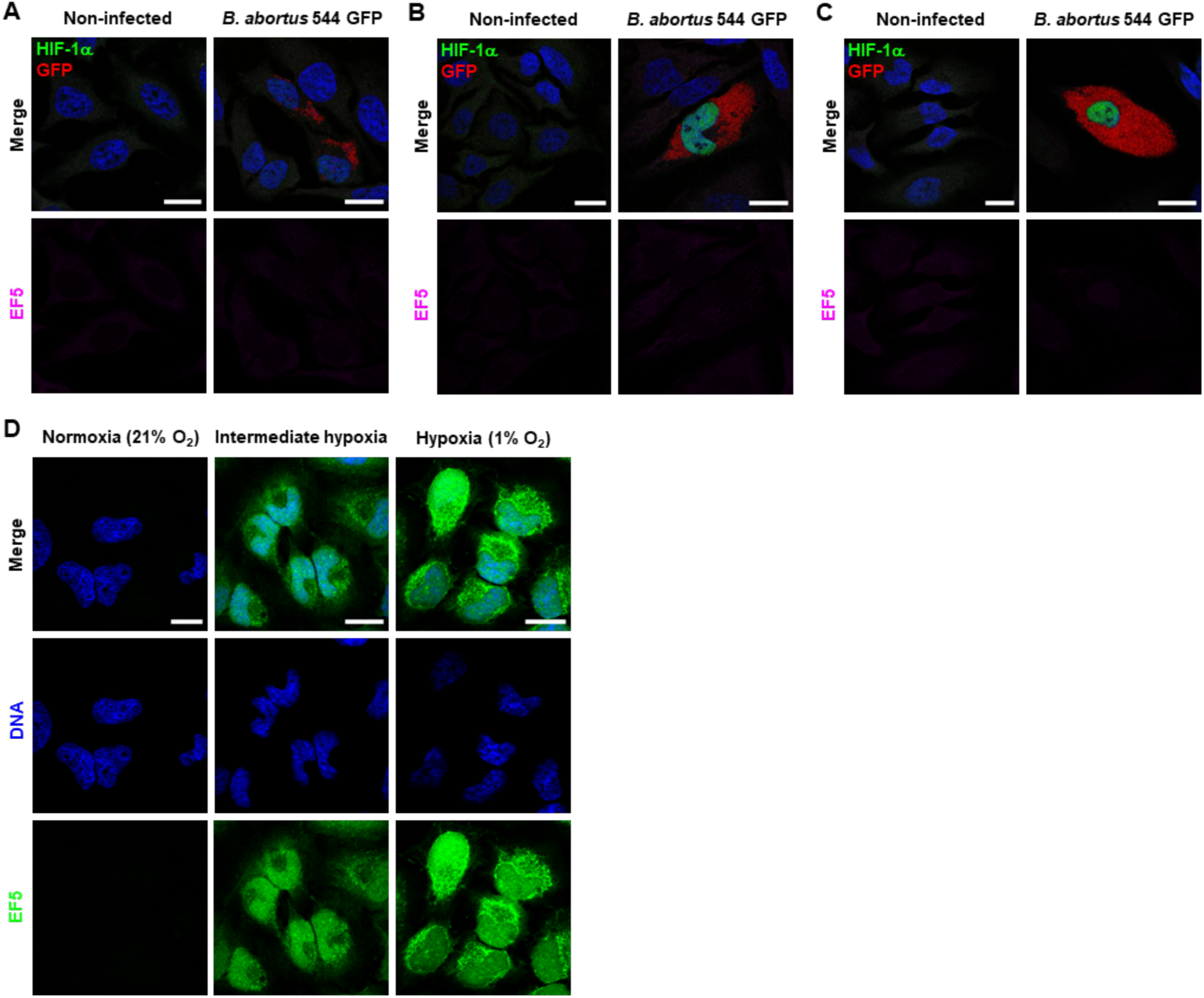
*B. abortus* induces HIF-1α stabilisation in a hypoxia-independent manner. A., B., C. Representative confocal micrographs of HeLa cells infected or not with *B. abortus* 544 GFP (Red) for 24 h (A), 48 h (B), and 72 h (C), treated with 150 μM of the EF5 compound for 3 h before analysis, then fixed and immunostained for EF5 (Anti-EF5 Cy5 conjugate – Magenta) and HIF-1α (Alexa 568 – Green). DNA was stained with Hoechst 33258 (Blue). Scale bars: 20 μm. D. Representative confocal micrographs of HeLa cells treated with 150 μM of the EF5 compound, exposed to normoxia (21 % O_2_), hypoxia (1 % O_2_) or an intermediate hypoxia (between 21 % and 1 % O_2_) for 3 h, then fixed and immunostained for EF5 (Anti-EF5 Cy5 conjugate – Green). DNA was stained with Hoechst 33258 (Blue). Scale bars: 20 μm.

**Figure S5.**
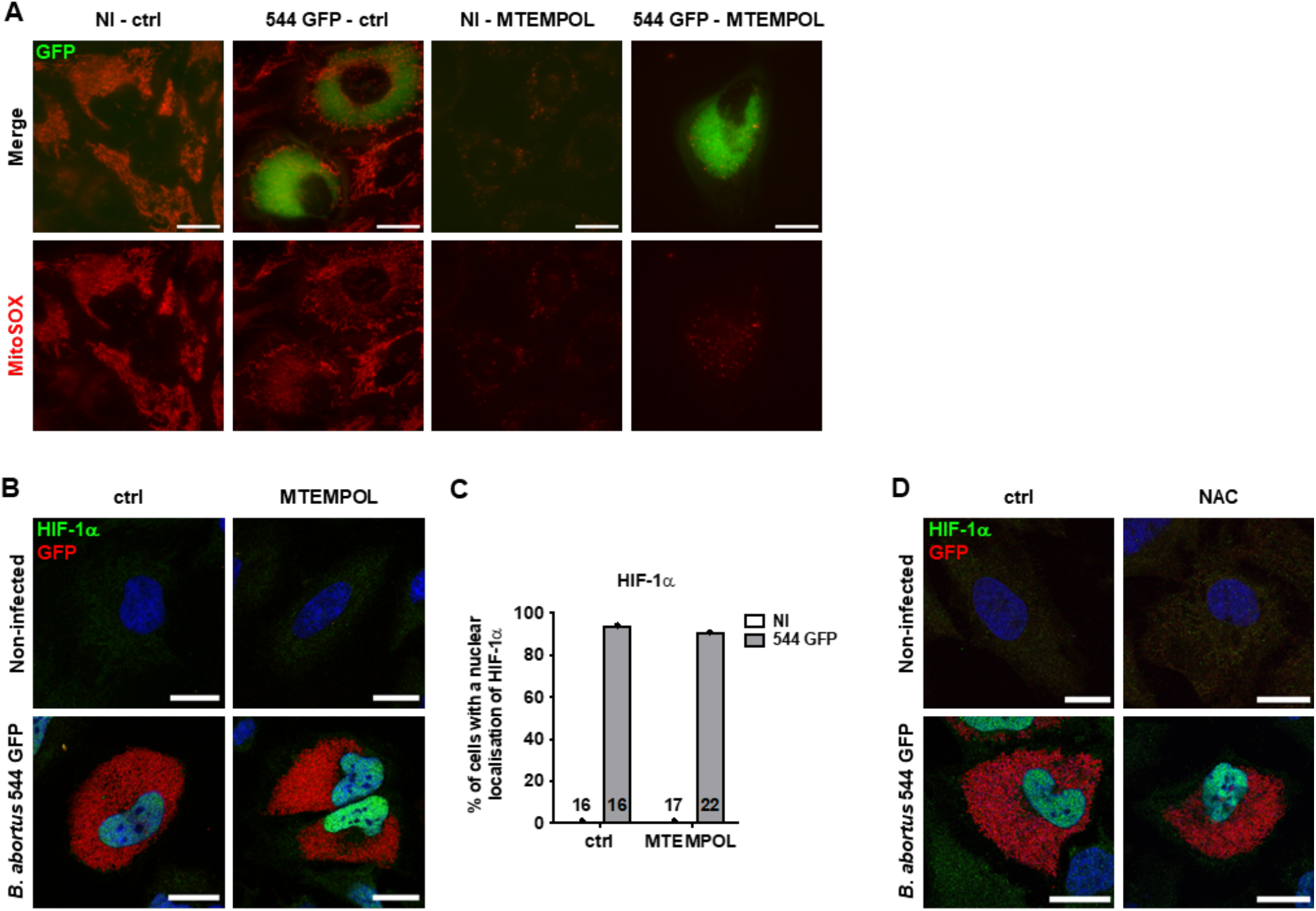
*B. abortus-induced* HIF-1α stabilisation is not mediated by mitochondrial ROS in HeLa cells. A. Representative wide-field micrographs of HeLa cells infected or not (NI) with *B. abortus* 544 GFP (Green) for 48 h, treated or not (ctrl) with 10 μM MTEMPOL for 24 h before analysis, and then stained with 0.5 μM MitoSOX™ fluorescent probe (Red) for 30 min. Samples were observed under live-imaging conditions with the Nikon Eclipse Ti2 inverted epifluorescence microscope. Scale bars: 20 μm. B. Representative confocal micrographs of HeLa cells infected or not (NI) with *B. abortus* 544 GFP (Red) for 48 h, treated or not (ctrl) with 10 μM MTEMPOL for 24 h before analysis, and then fixed and immunostained for HIF-1α (Alexa 568 – Green). DNA was stained with Hoechst 33258 (Blue). Scale bars: 20 μm. C. Quantification of the percentages of cells positive for a nuclear localisation of HIF-1α from HeLa cells infected or not (NI) with *B. abortus* 544 GFP and treated or not (ctrl) with 10 μM MTEMPOL for 24 h before analysis from micrographs shown in (B). Data are presented as mean from n=1 experiment (the numbers indicated in the columns represent the number of cells analysed per condition). D. Representative confocal micrographs of HeLa cells infected or not (NI) with *B. abortus* 544 GFP (Red) for 48 h, treated or not (ctrl) with 5 mM NAC for 24 h before analysis, and then fixed and immunostained for HIF-1α (Alexa 568 – Green). DNA was stained with Hoechst 33258 (Blue). Scale bars: 20 μm.

**Figure S6.**
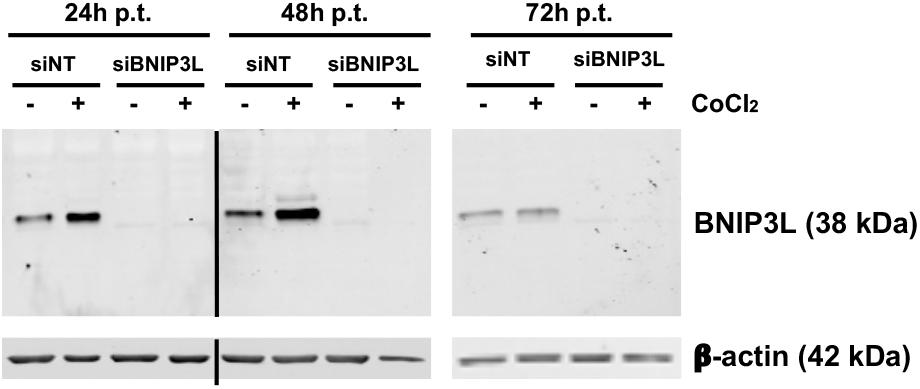
BNIP3L depletion efficiency in HeLa cells using a siRNA SMARTpool approach. Western-blot analysis of BNIP3L abundance in HeLa cells transfected with a non-targeting siRNA pool (siNT – 40 nM) or a BNIP3L siRNA SMARTpool (siBNIP3L – 40 nM) for 24 h, then left for the indicated times post-transfection (p.t.) and treated (+) or not (-) with 100 μM CoCl_2_ for 16 h before analysis. The abundance of β-actin was used as a loading control. Black bars indicate a crop in the membrane.

**Figure S7.**
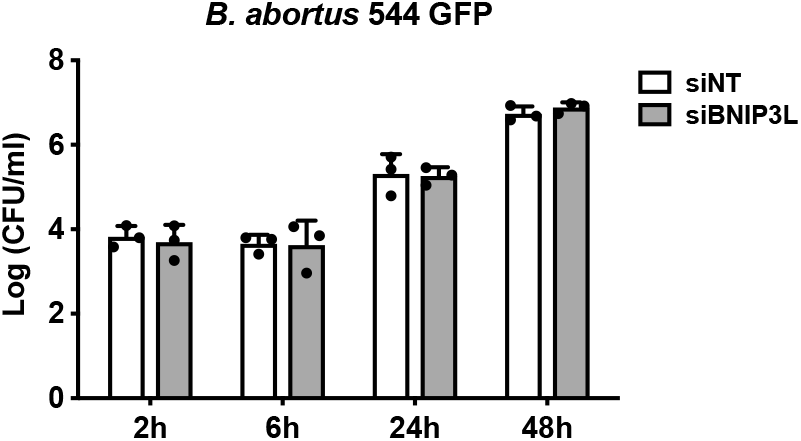
BNIP3L depletion does not impair *B. abortus* intracellular replication in HeLa cells. CFU assay expressing Log (CFU/ml) from HeLa cells transfected with a non-targeting siRNA pool (siNT – 40 nM) or a BNIP3L siRNA SMARTpool (siBNIP3L – 40 nM), then infected with *B. abortus* 544 GFP for the indicated times. Data are presented as mean ±SD from n=3 independent experiments; Statistical analyses were performed using a two-way ANOVA followed by a Šidàk’s multiple comparisons test; no significant differences were found.

## REFERENCES

Agarwal S & Muqit MMK (2022) PTEN-induced kinase 1 (PINK1) and Parkin: Unlocking a mitochondrial quality control pathway linked to Parkinson’s disease. Curr Opin Neurobiol 72: 111–119

Allen GFG, Toth R, James J & Ganley IG (2013) Loss of iron triggers PINK1/Parkin-independent mitophagy. EMBO Rep 14: 1127–1135

Atluri VL, Xavier MN, De Jong MF, Den Hartigh AB & Tsolis RM (2011) Interactions of the human pathogenic Brucella species with their hosts. Annu Rev Microbiol 65: 523–541

von Bargen K, Gorvel JP & Salcedo SP (2012) Internal affairs: Investigating the Brucella intracellular lifestyle. FEMS Microbiol Rev 36: 533–562

Bell EL, Klimova TA, Eisenbart J, Schumacker PT & Chandel NS (2007) Mitochondrial Reactive Oxygen Species Trigger Hypoxia-Inducible Factor-Dependent Extension of the Replicative Life Span during Hypoxia. Mol Cell Biol 27: 5737–5745

Borghesan E, Smith EP, Myeni S, Binder K, Knodler LA & Celli J (2021) A Brucella effector modulates the Arf6-Rab8a GTPase cascade to promote intravacuolar replication. EMBO J 40: 1–23

Boschiroli ML, Ouahrani-Bettache S, Foulongne V, Michaux-Charachon S, Bourg G, Allardet-Servent A, Cazevieille C, Liautard JP, Ramuz M & O’Callaghan D (2002) The Brucella suis virB operon is induced intracellularly in macrophages. PNAS 99: 1544–1549

Boutry M & Kim PK (2021) ORP1L mediated PI(4)P signaling at ER-lysosome-mitochondrion three-way contact contributes to mitochondrial division. Nat Commun 12: 1–18

Byndloss MX, Tsai AY, Walker GT, Miller CN, Young BM, English BC, Seyffert N, Kerrinnes T, de Jong MF, Atluri VL, et al (2019) Brucella abortus Infection of Placental Trophoblasts Triggers Endoplasmic Reticulum Stress-Mediated Cell Death and Fetal Loss via Type IV Secretion System-Dependent Activation of CHOP. MBio 10: 1–12

Carvalho F, Spier A, Chaze T, Matondo M & Cossart P (2020) Listeria monocytogenes Exploits Mitochondrial Contact Site and Cristae Organizing System Complex Subunit Mic10 To. MBio 11: 1–17

Celli J, de Chastellier C, Franchini D-M, Pizarro-Cerda J, Moreno E & Gorvel J-P (2003) *Brucella* Evades Macrophage Killing via VirB-dependent Sustained Interactions with the Endoplasmic Reticulum. J Exp Med 198: 545–556

Celli J, Salcedo SP & Gorvel J-P (2005) Brucella coopts the small GTPase Sar1 for intracellular replication. Proc Natl Acad Sci 102: 1673–1678

Chowdhury SR, Reimer A, Sharan M, Kozjak-Pavlovic V, Eulalio A, Prusty BK, Fraunholz M, Karunakaran K & Rudel T (2017) Chlamydia preserves the mitochondrial network necessary for replication via microRNA-dependent inhibition of fission. J Cell Biol 216: 1071–1089

Cloeckaert A, De Wergifosse P, Dubray G & Limet JN (1990) Identification of seven surface-exposed Brucella outer membrane proteins by use of monoclonal antibodies: Immunogold labeling for electron microscopy and enzyme-linked immunosorbent assay. Infect Immun 58: 3980–3987

Conway JRW, Warren SC, Herrmann D, Murphy KJ, Cazet AS, Vennin C, Shearer RF, Killen MJ, Magenau A, Mélénec P, et al (2018) Intravital Imaging to Monitor Therapeutic Response in Moving Hypoxic Regions Resistant to PI3K Pathway Targeting in Pancreatic Cancer. Cell Rep 23: 3312–3326

Copin R, Vitry MA, Hanot Mambres D, Machelart A, de Trez C, Vanderwinden JM, Magez S, Akira S, Ryffel B, Carlier Y, et al (2012) In situ microscopy analysis reveals local innate immune response developed around Brucella infected cells in resistant and susceptible mice. PLoS Pathog 8

Dai ZJ, Gao J, Ma X Bin, Yan K, Liu XX, Kang HF, Ji ZZ, Guan HT & Wang XJ (2012) Up-regulation of hypoxia inducible factor-1α by cobalt chloride correlates with proliferation and apoptosis in PC-2 cells. J Exp Clin Cancer Res 31: 28

Daskalaki I, Gkikas I & Tavernarakis N (2018) Hypoxia and selective autophagy in cancer development and therapy. Front Cell Dev Biol 6: 1–22

Dikic I & Elazar Z (2018) Mechanism and medical implications of mammalian autophagy. Nat Rev Mol Cell Biol 19: 349–364

Escoll P, Song OR, Viana F, Steiner B, Lagache T, Olivo-Marin JC, Impens F, Brodin P, Hilbi H & Buchrieser C (2017) Legionella pneumophila Modulates Mitochondrial Dynamics to Trigger Metabolic Repurposing of Infected Macrophages. Cell Host Microbe 22: 302–316.e7

Fine-Coulson K, Giguère S, Quinn FD & Reaves BJ (2015) Infection of A549 human type II epithelial cells with Mycobacterium tuberculosis induces changes in mitochondrial morphology, distribution and mass that are dependent on the early secreted antigen, ESAT-6. Microbes Infect 17: 689–697

Gomes MTR, Guimarães ES, Marinho F V., Macedo I, Aguiar ERGR, Barber GN, Moraes-Vieira PMM, AlvesFilho JC & Oliveira SC (2021) STING regulates metabolic reprogramming in macrophages via HIF-1α during Brucella infection. PLoS Pathog 17: 1–25

González-Espinoza G, Arce-Gorvel V, Mémet S & Gorvel JP (2021) Brucella: Reservoirs and niches in animals and humans. Pathogens 10: 1–21

Guo C, Zhang YX, Wang T, Zhong ML, Yang ZH, Hao LJ, Chai R & Zhang S (2015) Intranasal deferoxamine attenuates synapse loss via up-regulating the P38/HIF-1α pathway on the brain of APP/PS1 transgenic mice. Front Aging Neurosci 7

Hailey DW, Rambold AS, Satpute-Krishnan P, Mitra K, Sougrat R, Kim PK & Lippincott-Schwartz J (2010) Mitochondria Supply Membranes for Autophagosome Biogenesis during Starvation. Cell 141: 656–667

Hamasaki M, Furuta N, Matsuda A, Nezu A, Yamamoto A, Fujita N, Oomori H, Noda T, Haraguchi T, Hiraoka Y, et al (2013) Autophagosomes form at ER-mitochondria contact sites. Nature 495: 389–393

Kabeya Y, Mizushima N, Ueno T, Yamamoto A, Kirisako T, Noda T, Kominami E, Ohsumi Y & Yoshimori T (2003) LC3, a mammalian homolog of yeast Apg8p, is localized in autophagosome membranes after processing. EMBO J 22: 4577

Kang D, Kirienkoa DR, Webster P, Fisher AL & Kirienko N V. (2018) Pyoverdine, a siderophore from Pseudomonas aeruginosa, translocates into C. elegans, removes iron, and activates a distinct host response. Virulence 9: 804–817

Ke Y, Wang Y, Li W & Chen Z (2015) Type IV secretion system of Brucella spp. and its effectors. Front Cell Infect Microbiol 5: 1–10

Lee P, Chandel NS & Simon MC (2020) Cellular adaptation to hypoxia through hypoxia inducible factors and beyond. Nat Rev Mol Cell Biol 21: 268–283

Li X, Wu K, Zeng S, Zhao F, Fan J, Li Z, Yi L, Ding H, Zhao M, Fan S, et al (2021) Viral infection modulates mitochondrial function. Int J Mol Sci 22: 1–15

Liu Z, Reba S, Chen WD, Porwal SK, Boom WH, Petersen RB, Rojas R, Viswanathan R & Devireddy L (2014) Regulation of mammalian siderophore 2,5-DHBA in the innate immune response to infection. J Exp Med 211: 1197–1213

Lobet E, Letesson JJ & Arnould T (2015) Mitochondria: A target for bacteria. Biochem Pharmacol 94: 173–185

Lobet E, Willemart K, Ninane N, Demazy C, Sedzicki J, Lelubre C, De Bolle X, Renard P, Raes M, Dehio C, et al (2018) Mitochondrial fragmentation affects neither the sensitivity to TNFα-induced apoptosis of Brucella-infected cells nor the intracellular replication of the bacteria. Sci Rep 8: 1–17

Ma Z, Deng X, Li R, Hu R, Miao Y, Xu Y, Zheng W, Yi J, Wang Z, Wang Y, et al (2022) Crosstalk of Brucella abortus nucleomodulin BspG and host DNA replication process/mitochondrial respiratory pathway promote anti-apoptosis and infection. Vet Microbiol 268: 109414

Ma Z, Li R, Hu R, Deng X, Xu Y, Zheng W, Yi J, Wang Y & Chen C (2020) Brucella abortus BspJ Is a Nucleomodulin That Inhibits Macrophage Apoptosis and Promotes Intracellular Survival of Brucella. Front Microbiol 11: 1–16

Mahla RS, Kumar A, Tutill HJ, Krishnaji ST, Sathyamoorthy B, Noursadeghi M, Breuer J, Pandey AK & Kumar H (2021) NIX-mediated mitophagy regulate metabolic reprogramming in phagocytic cells during mycobacterial infection. Tuberculosis 126: 102046

Medeiros TC, Mehra C & Pernas L (2021) Contact and competition between mitochondria and microbes. Curr Opin Microbiol 63: 189–194

Miller CN, Smith EP, Cundiff JA, Knodler LA, Bailey J, Lupashin V & Celli J (2018) A Brucella Type IV effector targets the COG tethering complex to remodel host secretory traffic and promote intracellular replication. Cell Host Microbe 22: 317–329

Mills EL, Kelly B & O’Neill LAJ (2017) Mitochondria are the powerhouses of immunity. Nat Immunol 18: 488–498

Missiroli S, Genovese I, Perrone M, Vezzani B, Vitto VAM & Giorgi C (2020) The role of mitochondria in inflammation: From cancer to neurodegenerative disorders. J Clin Med 9

Mukhopadhyay P, Rajesh M, Yoshihiro K, Haskó G & Pacher P (2007) Simple quantitative detection of mitochondrial superoxide production in live cells. Biochem Biophys Res Commun 358: 203–208

Narendra D, Tanaka A, Suen DF & Youle RJ (2008) Parkin is recruited selectively to impaired mitochondria and promotes their autophagy. J Cell Biol 183: 795–803

Onishi M & Okamoto K (2021) Mitochondrial clearance: mechanisms and roles in cellular fitness. FEBS Lett 595: 1239–1263

Onishi M, Yamano K, Sato M, Matsuda N & Okamoto K (2021) Molecular mechanisms and physiological functions of mitophagy. EMBO J 40: 1–27

Parent MA, Bellaire BH, Murphy EA, Roop RM, Elzer PH & Baldwin CL (2002) Brucella abortus siderophore 2,3-dihydroxybenzoic acid (DHBA) facilitates intracellular survival of the bacteria. Microb Pathog 32: 239–248

Roba A, Carlier E, Godessart P, Naili C & De Bolle X (2022) Histidine auxotroph mutant is defective for cell separation and allows the identification of crucial factors for cell division in Brucella abortus. Mol Microbiol

Roop II RM (2012) Metal acquisition and virulence in Brucella. Anim Heal Res Rev 13: 1–7

Roset MS, Alefantis TG, DelVecchio VG & Briones G (2017) Iron-dependent reconfiguration of the proteome underlies the intracellular lifestyle of Brucella abortus. Sci Rep 7: 10637

Sandoval H, Thiagarajan P, Dasgupta SK, Schumacher A, Prchal T, Chen M & Wang J (2008) Essential role for Nix in autopagic maturation of erythroid cells. Nature 454: 232–235

Sassera D, Beninati T, Bandi C, Bouman EAP, Sacchi L, Fabbi M & Lo N (2006) ‘Candidatus Midichloria mitochondrii’, an endosymbiont of the Ixodes ricinus with a unique intramitochondrial lifestyle. Int J Syst Evol Microbiol 56: 2535–2540

Schofield CJ & Ratcliffe PJ (2004) Oxygen sensing by HIF hydroxylases. Nat Rev Mol Cell Biol 5: 343–354

Sedzicki J, Tschon T, Low SH, Willemart K, Goldie KN, Letesson JJ, Stahlberg H & Dehio C (2018) 3D correlative electron microscopy reveals continuity of Brucella-containing vacuoles with the endoplasmic reticulum. J Cell Sci 131

Shi J, Yu J, Zhang Y, Wu L, Dong S, Wu L, Wu L, Du S, Zhang Y & Ma D (2019) PI3K/Akt pathway-mediated HO-1 induction regulates mitochondrial quality control and attenuates endotoxin-induced acute lung injury. Lab Investig 99: 1795–1809

Sirianni A, Krokowski S, Lobato-Márquez D, Buranyi S, Pfanzelter J, Galea D, Willis A, Culley S, Henriques R, Larrouy-Maumus G, et al (2016) Mitochondria mediate septin cage assembly to promote autophagy of Shigella. EMBO Rep 17: 1029–1043

Smith EP, Cotto-Rosario A, Borghesan E, Held K, Miller CN & Celli J (2020) Epistatic Interplay between Type IV Secretion Effectors Engages the Small GTPase Rab2 in the Brucella Intracellular Cycle. MBio 11: 1–14

Smith JA, Khan M, Magnani DD, Harms JS, Durward M, Radhakrishnan GK, Liu YP & Splitter GA (2013) Brucella Induces an Unfolded Protein Response via TcpB That Supports Intracellular Replication in Macrophages. PLoS Pathog 9: 1–12

Spier A, Stavru F & Cossart P (2019) Interaction between Intracellular Bacterial Pathogens and Host Cell Mitochondria. Microbiol Spectr 7: 1–11

Starr T, Child R, Wehrly TD, Hansen B, Hwang S, López-Otin C, Virgin HW & Celli J (2012) Selective subversion of autophagy complexes facilitates completion of the Brucella intracellular cycle. Cell Host Microbe 11: 33–45

Starr T, Ng TW, Wehrly TD, Knodler LA & Celli J (2008) Brucella intracellular replication requires trafficking through the late endosomal/lysosomal compartment. Traffic 9: 678–694

Stavru F, Riemer J, Jex A & Sassera D (2020) When bacteria meet mitochondria: The strange case of the tick symbiont Midichloria mitochondrii†. Cell Microbiol 22: 1–9

Szabadkai G, Simoni AM, Chami M, Wieckowski MR, Youle RJ & Rizzuto R (2004) Drp-1-Dependent Division of the Mitochondrial Network Blocks Intraorganellar Calcium Waves and Protects against Calcium-Mediated Apoptosis. Mol Cell 16: 59–68

De Vos KJ & Sheetz MP (2007) Visualization and Quantification of Mitochondrial Dynamics in Living Animal Cells. Methods Cell Biol 80: 627–682

Wang Y, Nartiss Y, Steipe B, McQuibban GA & Kim PK (2012) ROS-induced mitochondrial depolarization initiates PARK2/PARKIN-dependent mitochondrial degradation by autophagy. Autophagy 8: 1462–1476

Zachari M & Ktistakis NT (2020) Mammalian Mitophagosome Formation: A Focus on the Early Signals and Steps. Front Cell Dev Biol 8: 1–11

Zhang Y, Yao Y, Qiu X, Wang G, Hu Z, Chen S, Wu Z, Yuan N, Gao H, Wang J, et al (2019) Listeria hijacks host mitophagy through an novel mitophagy receptor to evade killing. Nat Immunol 20: 433–446

